# Cell cycle-driven transcriptome maturation confers multilineage competence to cardiopharyngeal progenitors

**DOI:** 10.1101/2024.07.23.604718

**Authors:** Yelena Y. Bernadskaya, Ariel Kuan, Andreas Tjärnberg, Jonas Brandenburg, Ping Zheng, Keira Wiechecki, Nicole Kaplan, Margaux Failla, Maria Bikou, Oliver Madilian, Wei Wang, Lionel Christiaen

## Abstract

During development, stem and progenitor cells divide and transition through germ layer- and lineage-specific multipotent states to generate the diverse cell types that compose an animal. Defined changes in biomolecular composition underlie the progressive loss of potency and acquisition of lineage-specific characteristics. For example, multipotent cardiopharyngeal progenitors display multilineage transcriptional priming, whereby both the cardiac and pharyngeal muscle programs are partially active and coexist in the same progenitor cells, while their daughter cells engage in a cardiac or pharyngeal muscle differentiation path only after cell division. Here, using the tunicate Ciona, we studied the acquisition of multilineage competence and the coupling between fate decisions and cell cycle progression. We showed that multipotent cardiopharyngeal progenitors acquire the competence to produce distinct *Tbx1/10* (+) and (−) daughter cells shortly before mitosis, which is necessary for *Tbx1/10* activation. By combining transgene-based sample barcoding with single cell RNA-seq (scRNA-seq), we uncovered transcriptome-wide dynamics in migrating cardiopharyngeal progenitors as cells progress through G1, S and G2 phases. We termed this process “transcriptome maturation”, and identified candidate “mature genes”, including the Rho GAP-coding gene *Depdc1*, which peak in late G2. Functional assays indicated that transcriptome maturation fosters cardiopharyngeal competence, in part through multilineage priming and proper oriented and asymmetric division that influences subsequent fate decisions, illustrating the concept of “behavioral competence”. Both classic feedforward circuits and coupling with cell cycle progression drive transcriptome maturation, uncovering distinct levels of coupling between cell cycle progression and fateful molecular transitions. We propose that coupling competence and fate decision with the G2 and G1 phases, respectively, ensures the timely deployment of lineage-specific programs.

## Introduction

Complex animals are characterized by dozens to hundreds of distinct cell types that emerge during embryogenesis, and post-embryonic development^1–3^. During this process, the developmental potential of successive generations of pluri- and multipotent progenitor cells is progressively restricted, while they acquire the competence to produce a few differentiated cell types^4^. Changes in the biomolecular composition of cells underlie these developmental transitions, and differential transcriptional activity governing transcriptome dynamics is an established driver of fateful molecular transitions during development^5,6^.

While certain differentiated cells, such as neurons and striated muscles, are typically post-mitotic, multipotent progenitor cells must divide to express their full potential and produce a variety of cell lineages. The molecular machinery driving cell cycle progression and division is well characterized, and largely conserved across developmental and phylogenetic lineages^7–9^. How the cell cycle interfaces with fate choices during development remains debated, and is likely to be highly variable. On one hand, there is evidence that fate choices can occur independently of cell cycle progression ^10,11^, to the point that so-called “cell cycle genes” are often *regressed out* of single cell genomics analyses aimed at charting development decisions^12,13^. On the other hand, the G1 phase of the cell cycle tends to increase the propensity for mammalian stem cells to activate fateful determinants and engage along a certain developmental path^14–17^, while certain neuroblasts in *Drosophila* ^18,19,20^, and early blastomeres in ascidian and *C. elegans* embryos appear to change fate with every division. Conversely, cell fate choices have been shown to directly impact the cell cycle in a variety of organisms. For example, developmental regulation of the phosphatase Cdc25 contributes to coordinating mitotic patterns in the early fly embryo^21–24^, and differential beta-catenin activity, or *Cdc25* or *Cdkn1* expression accounts for blastomere and lineage-specific timing of cell division in ascidians^25–27^.

The concept of multipotent cardiopharyngeal progenitors emerged as a compelling paradigm to account for the shared cardiac and craniofacial congenital defects observed in various condition, such as the Di George/22q11 deletion syndrome, which is often caused by large deletions that remove a copy of the *TBX1* gene^28–30^. This T-box transcription factor-coding gene is expressed early in progenitor cells for both the anterior second heart field and the branchiomeric skeletal muscle, both of which require its function for proper heart and head muscle development^31–33^. Notably, ascidian tunicates of the *Ciona* genus possess a well-defined cardiopharyngeal lineage, which emerges from *Mesp*+ mesodermal progenitors as is the case in mammals, and produces a *Gaca4/5/6*-positive and *Tbx1*-negative first heart lineage, as well as the hallmark *Tbx1/10*+ multipotent progenitors for the second heart and pharyngeal muscle lineages^34,35^ (**Figure 1A**). The ascidian cardiopharyngeal lineage develops in a highly stereotyped fashion, providing single cell resolution view of the interplay between cardiopharyngeal multipotency, cell divisions and heart vs. pharyngeal muscle fate choices.

Here, we leveraged the unique features and experimental amenability of the *Ciona* embryo to study the molecular mechanisms underlying the acquisition of multilineage competence in cardiopharyngeal progenitors. We describe their transcriptome maturation, identifying a mature state that confers competence to divide in an oriented and asymmetric fashion, and produce distinct *Tbx1/10*(+) second multipotent cardiopharyngeal progenitors and *Tbx1/10*(−) first heart precursor cells. We characterized the regulation of multipotent progenitor maturation, and identified distinct levels of coupling between cell cycle progression and progenitor maturation and fate choices.

## Results

### Mitosis is necessary but not sufficient for *Tbx1/10* activation in cardiopharyngeal progenitors

The conserved cardiopharyngeal determinant *Tbx1/10* is activated after division of multipotent cardiopharyngeal progenitors (aka Trunk Ventral Cells, TVCs), following collective migration^36,37^. These oriented and unequal cleavages are coupled with asymmetric Fibroblast Growth Factor (FGF)-Microtubule Associated Protein Kinase (MAPK) signaling^37^, which positions the first heart progenitors medially and the *Tbx1/10*+ second-generation multipotent progenitors (aka second trunk ventral cells, STVCs) laterally (Figure 1A-D’). As is the case in vertebrates, Tbx1/10 promotes pharyngeal muscle specification, in part by antagonizing the cardiac fate^36–39^. It is thus essential that *Tbx1/10* be activated after division of multipotent progenitors, to allow for the emergence of the first heart lineage, which produces the majority of cardiomyocytes in Ciona^35^.

**Figure 1.**
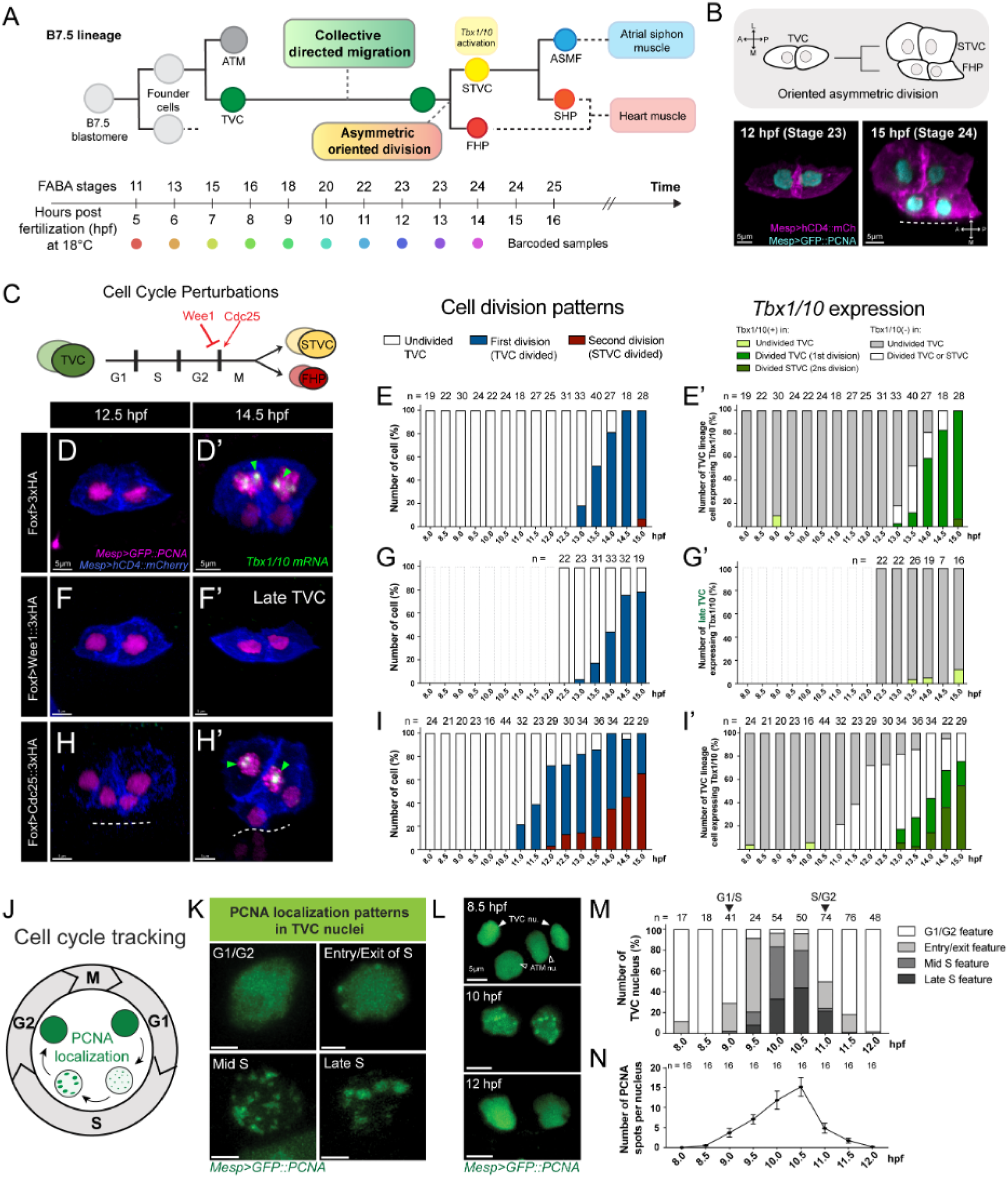
Mitosis is necessary but not sufficient for Tbx1/10 activation in cardiopharyngeal progenitors. **Progression through the progenitor cell cycle promotes expression of cardiopharyngeal fate determinants** **A.** (Top) Developmental trajectory of the cardiopharyngeal lineage in basal chordate *Ciona robusta*. A B7.5 lineage from one side of the embryo is shown, where a single B7.5 cell gives rise to a pair of founder cells, producing a pair of TVCs and ATMs. (Bottom) Correspondence of FABA stages, post-fertilization developmental time points (18°C), and color codes of sc-RNAseq barcodes are shown. TVC, trunk ventral cell; ATM, anterior tail muscle; STVC, second trunk ventral cell; FHP, first heart precursor; ASMF, atrial siphon muscle founder cells; SHP, second heart precursor. **B.** (Top) Schematic diagram of asymmetric oriented division of TVCs. (Bottom) Confocal images of before (left, 12 hpf) and after (right, 15 hpf) TVC division. Cyan: nuclei (NLS::LacZ); Magenta: cell membranes (hCD4::mCherry). Dashed line represents the embryo midline. M, medial; L, lateral. Scale bar = 5mm. **C.** Schematic of TVC cell cycle stages and genetic perturbations of mitotic entry. **D-I’**. Cell division patterns (E, G, I) and Tbx1/10 expression (E’, G’, I’) during cell cycle perturbations. Control TVC division (D-E, 3’HA), inhibition of TVC mitotic entry (F-G, Wee1::3’HA), and induction of TVC mitotic entry (H-I, Cdc25::3’HA) conditions are examined from 8 to 15 hpf. Perforated bars in G and G’ indicate timepoints not analyzed. Magenta: nuclei (GFP::PCNA); Blue: cell membranes (hCD4::mCherry); Green arrowhead: Tbx1/10 mRNA, Scale bar = 5mm. **J.** Schematic of variability of PCNA puncta patterns in the TVC nuclei associated with progression throughout the cell cycle. **K.** Confocal images of PCNA puncta distribution in individual TVC nuclei at different cell cycle stages. Green: GFP::PCNA. Scale bar = 2.5mm. **L-N.** Determination of S phase of TVC using PCNA. GFP::PCNA are visualized in the B7.5-lineage under the Mesp enhancer at 8-12 hpf. Representative confocal images showing the G1, S, and G2 stages of TVC at 8, 10, and 12 hpf (L). Green: GFP::PCNA. Scale bar = 5mm. Developmental distribution of four PCNA localization patterns (M). Quantification of PCNA spots per nucleus across developmental stages (N).

We previously showed that division of multipotent cardiopharyngeal progenitors is necessary for *Tbx1/10* activation, which is not detected following lineage-specific inhibition of either G1-S or G2-M transition by misexpression of Cdkn1 or Wee1, respectively^37^. Here we confirmed these results, and determined that *Tbx1/10* expression does not recover in division-inhibited cardiopharyngeal progenitors (**Figure 1C-G**), unlike *Ebf*, the pharyngeal muscle determinant that is merely delayed by mitosis inhibition^37^. When only one of the two cardiopharyngeal progenitors inherited the *Foxf*-enhancer-driven Wee1-expressing plasmid, which typically occurs in ∼40% of embryos due to mosaic incorporation of electroporated transgenes^40^, only the unaffected cell induced *Tbx1/10* expression, indicating that cell division is required cell-autonomously (Figure 1F-G’ and Figure 1-S1). Mitosis is thus necessary for *Tbx1/10* activation in cardiopharyngeal progenitors.

Next, we sought to induce precocious division of cardiopharyngeal progenitors and test whether mitosis also sufices to triger *Tbx1/10* expression. To this aim, we reused the minimal *Foxf* cardiopharyngeal enhancer^41^ to misexpress Cdc25, a conserved phosphatase that antagonizes the Wee1 kinase and promotes entry into mitosis by dephosphorylating the mitotic Cyclin-Dependent Kinase (CDK)^42^. Consistent with a stereotyped and synchronous developmental progression, control cardiopharyngeal progenitors typically divide between 13 and 14 hours post-fertilization (hpf) at 18°C (FABA stage 23 and 24^43^) (**Figure 1A, D-E**). By contrast, Cdc25 over-expression (Cdc25^OE^) caused progenitor cells to divide as early as 11 hpf in ∼20% of embryos, completing mitosis and proceeding to the second division approximately 2 hours earlier than the equivalent cells in control embryos (H-I), albeit with substantial defects in orientation and asymmetry (**Figure 1-S2**).

Having established Cdc25^OE^ as a reliable method to induce precocious division of multipotent cardiopharyngeal progenitors, we leveraged this perturbation to assay the sufficiency of mitosis for *Tbx1/10* activation (**Figure 1H-I’**). In controls, the proportions of embryos with *Tbx1/10*+ cells become equivalent to the percentage of embryos with divided cells within approximately 30 minutes, suggesting that mitosis is rapidly followed by the onset of *Tbx1/10* transcription, as assayed by intronic probes that detect nascent transcripts^36^ (**Figure 1D’,E’**). By contrast, while Cdc25^OE^ caused precocious mitosis, *Tbx1/10* expression was first detected approximately 2 hours later, starting at around the same time as in control embryos (**Figure 1H-’, Figure 1-S3**). Careful examination of *Tbx1/10* expression in Cdc25^OE^ cells indicated a slight reduction in expression and a sustained pattern of lateral-expression after oriented divisions (**Figure 1-S4**). Taken together, these data indicate that mitosis is necessary but not suficient to triger *Tbx1/10* expression in the cardiopharyngeal lineage.

To better understand cell cycle progression and the Cdc25^OE^ phenotype, we used the PCNA::GFP marker, a helicase that localizes to DNA replication forks, forming conspicuous nuclear dots during S-phase^25^ (**Figure 1J-N**). PCNA::GFP nuclear dots tend to increase in size and become less numerous as replication forks coalesce and genome duplication approaches completion^44^ (**Figure 1J,K**), a feature that we leveraged to evaluate S-phase progression, by quantifying the number and size distribution of PCNA spots in time series of fixed embryos (**Figure 1L-N**). These analyses indicated that multipotent progenitors enter S-phase at approximately stage 18 (9 hpf at 18°C) and exit at stage 22 (∼11 hpf), albeit with occasional asynchrony between the leader and trailer cells (**Figure 1-S5**). Using this approach to examine cell cycle progression following Cdc25^OE^, we determined that the G1/S transition was shifted only approximately 30 min compared to controls, which could be explained by the potential of Cdc25 in facilitating G1/S transition. As expected however, the main effect of Cdc25^OE^ was to trigger precocious mitosis in cells that had presumably completed S-phase and entered G2, thus reducing the duration of G2 from ∼2 hours to less than 30 minutes (**Figure 1-S5**).

### Single cell developmental trajectories reveal transcriptome maturation in multipotent progenitors

Among various possible explanations, the lack of precocious *Tbx1/10* expression following forced division suggested that cardiopharyngeal progenitors become competent for mitosis-dependent *Tbx1/10* activation toward the end of interphase, once cells have completed directed collective migration and S-phase, and advanced into G2. Both cell-autonomous and cell-extrinsic factors determine competence and instruct fate decisions, including the transcription factor Hand-r/NoTrlc and FGF-MAPK signaling, which induce *Tbx1/10* activation in the cardiopharyngeal lineage^37^. However, these known regulators are already active in early cardiopharyngeal progenitors and unlikely to be limiting for early *Tbx1/10* activation following precocious cell division.

Focusing on cell-autonomous determinants of cardiopharyngeal competence, we harnessed single cell RNA-seq to gain insights into transcriptome dynamics as multipotent progenitors progress through the cell cycle, and migrate collectively, until they divide to produce first heart precursors (aka FHP) and *Tbx1/10*+ multipotent cardiopharyngeal progenitors (aka second trunk ventral cells, STVCs). We reasoned that FAC-sorting cardiopharyngeal lineage cells from embryos collected every hour through 10 time points encompassing the whole 7-hour interphase would leverage the natural variability between individual cells, and allow for the reconstruction of a developmental trajectory providing high-resolution insights into transcriptome dynamics.

We streamlined the experiment, and avoided technical batch effects, by developing a multiplexing approach to collect the entire dataset in one experiment. We created a library of 20 reporter constructs each containing a unique 9-nucleotide barcode in the 3’UTR of the *Mesp>GFP* reporter that labels the B7.5/cardiopharyngeal lineage (**Figures 2A, 2-S1**). We positioned the barcodes to optimize recovery by RNA sequencing, and used pairs of unique barcoded reporters for each individual sample. To obtain a ten time-points series, we took advantage of *en masse* electroporation of Ciona eggs to generate samples fertilized and transfected with pairs of unique barcoded reporters every hour, and collected all samples 14 hours after the first fertilization and electroporation. This approach yielded a whole 5 to 14 hpf time series encompassing stages 11 to 24/25, starting with late gastrula embryos containing *Mesp*+ naive mesodermal progenitors (aka founder cells), prior to the birth of multipotent cardiopharyngeal progenitors, and ending with pre-hatching larvae, which possess first heart precursors and second *Tbx1/10*+ multipotent progenitors after division (**Figures 1A, 2A**). Batches of embryos were individually dissociated and cell suspensions methanol-fixed and pooled for storage at −80°C. The sample was then rehydrated, immunolabeled with a fluorescent anti-GFP antibody for FAC-sorting of B7.5 lineage cells prior to loading onto a 10X Genomics Chromium controller.

This first experiment yielded 2,595 high-quality single cell transcriptomes where 500 to 70,000 reads detected 428 to 6,059 expressed genes in 98% of the cells (**Figure 2-S1A**). We processed single cell transcriptome data using standard methods incorporated in the Scanpy package^13^, in addition to our self-supervised graph-based denoising method DEWAKSS^45^, which led us to use 35 principal components for dimensionality reduction and k=18 nearest neighbors for denoising (**Figure 2-S1B,C**).

We specifically amplified and sequenced the barcode-containing regions of expressed transgenes in order to maximize recovery of sample barcodes in individual transcriptomes. The 20 SBC barcodes were added to the *Ciona robusca* reference genome as pseudogenes, and the raw sequencing data was mapped to the genome using 10x Genomics Cell Ranger to generate a UMI count matrix comprising the sample barcodes. We could assign sample barcodes to 50% of the single cell transcriptomes (1,261/2,517; **Figure 2-S1**E). Remarkably, the barcoded transcriptomes pertained to 6 related clusters separated from the unmarked clusters as illustrated on a projection of the multidimensional dataset in two dimensions using the UMAP algorithm (**Figures 2B**, **2-S1F,G**). Inspection of known markers *Mesp/MSGN1*, *Foxf, Gaca4/5/6* and *Myod/Myf5* confirmed that the clusters containing most barcoded transcriptomes correspond to B7.5 lineage cells (**Figures 2C,2-S1G-I**). The non-barcoded clusters thus reflect “contamination” by cells outside the *Mesp>GFP*-labeled B7.5 lineage during the FACS procedure, including - for example - *Foxf*+ trunk epidermal cells. This indicated that most B7.5 lineage cells were effectively barcoded, and enriched approximately 500 times in the sorted sample, since *Mesp>GFP*+ cells typically represent <0.1% of whole embryo cell suspensions^46^.

With the exception of SBC00 and SBC20, individual barcodes displayed high (>0.98) correlation coefficients between cotransfected pairs, and very low (<0.05) correlation with other barcodes electroporated independently (**Figure 2-S1D**). This indicated that our transgene-based barcoding and multiplexing strategy allowed us to process pooled samples and demultiplex datasets *in silico*, assigning a time point of origin to most individual single cell transcriptomes obtained from B7.5 lineage cells, albeit with a disparate representation of each sample (**Figure 2-S1E**).

Our lineage-specific time series lent itself to developmental trajectory analysis, as two dimensional UMAP projection clearly distinguished between the anterior tail muscle and cardiopharyngeal/trunk ventral cell trajectories, both stemming from shared origins in the naive *Mesp*+ mesoderm (aka founder cells; **Figure 1A, Figure 2B-D**). Pseudotime inference corroborated UMAP projection and was concordant with real time distributions, illustrating that temporal changes in transcriptome composition account for most of the variance (**Figure 2E-G**). Gene expression denoising and mapping cells onto lineage-specific pseudotemporal sequences provided a high resolution view of gene expression dynamics, as illustrated by the sequential activations of key regulators of cardiopharyngeal development *Mesp*, *Ets1/2*, *Foxf*, *Gaca4/5/6*, *Nk4/Nkx2.5* and *Hand1/2* (**Figure 2-S2A**).

Leveraging this high-resolution view of whole transcriptome dynamics, we identified clusters of genes displaying distinct (pseudo)temporal profiles of expression along the anterior tail muscle and cardiopharyngeal progenitor trajectories, separately (**Figure 2H-I**, **Figure 2-S2B**). We reasoned that seemingly continuous gene expression changes in fact reflect transitions between successive regulatory states^4^. We used a clustering approach to segment individual trajectories into significantly different states (**Figure 2H,I, 2-S2B**). In the primary tail muscle trajectory, 8 clusters of genes reflected transition through 3-to-4 regulatory states that seemingly stabilize as early as ∼9 hpf in a state marked by upregulation of cluster 6 genes, including *Myod/Myf5*, *Smyd1* and *Myl1* genes, and downregulation of cluster 0 and 2 genes, comprising naive *Mesp*+ mesoderm markers and primed cardiopharyngeal genes such as *Rgs21*, *Ccna*, *Zfp36L1/2*, which indicated differentiation toward a tail muscle cell type in post-mitotic cells (**Figure 2-S2B**). In the cardiopharyngeal progenitor trajectory, cells transitioned through 5 predicted regulatory states, each marked by a combination of relative expression for 8 clusters of genes. Notably, these predicted states do not appear to be stable, and could instead be considered as “transition states”^4^. For example, states 2 and 3 differ primarily by the relative dynamics of gene clusters 2 and 6, whereby cluster 2 appears to peak in state 3 after an expression onset during state 2, while cluster 6 genes peak in states 1 and 2 and become downregulated in state 3 (Figure 2I). Notably, gene clusters 0, 1 and 6 appear to be “off” and cluster 2 becomes downregulated in state 4, whereas cluster 4 and 5 peak toward the end of the cardiopharyngeal trajectory. State 4 comprises mostly cells collected from >11 hpf embryo, suggesting that it corresponds to the G2 phase, which sees the emergence of the cellular competence to divide and activate *Tbx1/10* (Figure 1). We thus considered regulatory state 4 as the “mature state”, characterized by low expression of gene clusters 0,1 and 6, downregulation of cluster 2 and peak expression of clusters 4 and 5.

**Figure 2.**
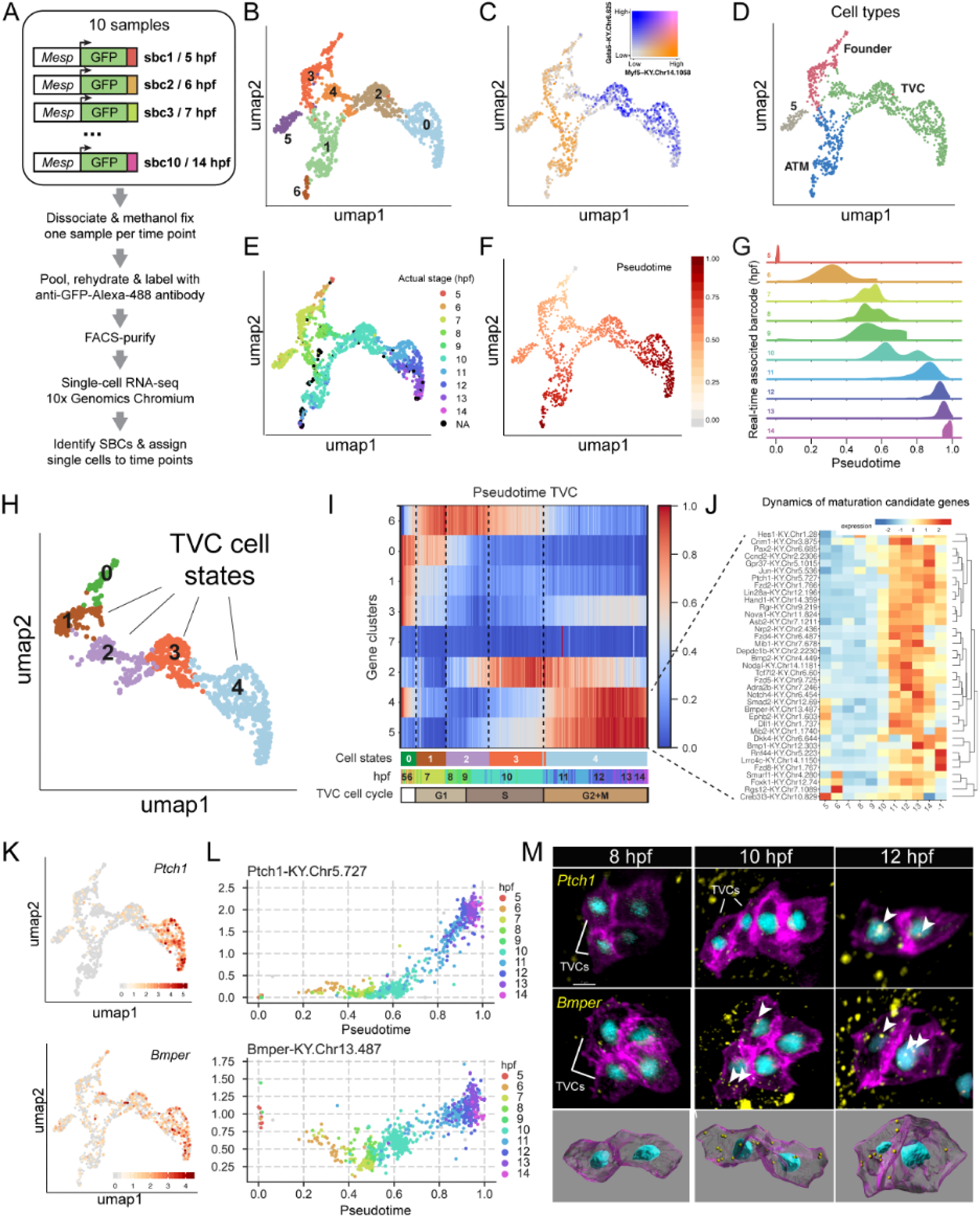
Single-cell developmental trajectories reveal transcriptome maturation in multipotent progenitors. **A.** scRNA-seq workflow outlining lineage-specific real-time barcoding strategy. **B.** Denoised Leiden clustering of anterior tail muscles and cardiopharyngeal trajectories **C.** Trajectory-specific expression of the TVC-specific Gata5 transcripts and founder and ATM-specific Myf5 transcripts. **D.** Assignment of cell type to trajectories based on lineage-specific gene expression. **E.** Reconstruction of cell trajectories using recovered real-time barcodes. **F.** Trajectory of pseudotime assignment in recovered cell trajectories. **G.** Distribution of real-time associated barcodes over pseudotime. **H.** Cell states of the TVC trajectory derived from agglomerative clustering. **I.** Differential expression of genes during TVC migration and subsequent cell division. Gene cluster dynamics show unique distributions that align with transition of proposed cell states. **J.** Dynamics of cluster 4 and cluster 5 genes selected for use in the targeted CRISPER screen. **K-M.** Validation of predicted dynamics of 2 cluster 5 genes, Ptch1 and BMPER. K. Expression of Ptch1 and BMPER in the cardiopharyngeal precursors. **L.** Dynamics in Ptch1 and BMPER expression levels as a function of pseudotime. Color code corresponds to barcodes assigned to developmental real time. **M.** Fluorescent in situ hybridization showing activation of Ptch1 and BMPER transcription in the cardiopharyngeal lineage. Nuclei are marked with NLS::LacZ (cyan), membranes are marked with hCD4::mCh (magenta), and transcripts (yellow) are indicated with arrowheads. Bottom row shows segmented TVCs with transcripts detected as yellow spots.

Importantly, dynamic genes comprised developmental regulators and cell biological effectors in addition to classic “cell cycle genes”, such as *Cdc25*, suggesting that the mature state does correspond to a developmentally significant regulatory state (**Figure 2I,J**). Finally, both published and new whole mount *in situ* hybridization assays corroborated the predicted expression dynamics for selected genes (**Figure 2K-M, Figure 2-S2C**). For example, the signaling molecule coding genes *Ptch1* and *Bmper*, were predicted to be activated specifically in the cardiopharyngeal progenitor lineage (**Figure 2K**) toward the end of the mature state (**Figure 2L**), which we confirmed by fluorescence *in situ* hybridization (Figure 2M).

### Generalization to the whole embryo

To expand the above proof of concept, we designed a library of barcoded reporters using the ubiquitously active *Ef1-alpha* driver, and repeated the time series collection between 5 and 14 hpf at 18°C followed by cell dissociation, pooling, methanol fixation, rehydration and single cell RNA-seq, but omitted the FACS step to profile the entire developmental sequence of the whole embryo in one experiment. We obtained >21,000 single cell transcriptomes altogether expressing >11,000 genes and distributed across 57 clusters (**Figure 3A**). We recovered temporal barcodes for 42.6% (8,974/21,087) of the cells, which is markedly lower than in our B7.5 lineage-focused experiment, because we did not FACS-purify transfected cells (**Figure 3B, 3-S1**). Nonetheless, all clusters contained barcoded transcriptomes, allowing us to map them back onto the developmental sequence, with the notable exception of the germline (cluster 54, **Figure 3**), which does not effectively transcribe electroporated transgenes^47^. Using a label transfer approach, we expanded the barcode-derived assignment to 86.6% of the dataset (**Figure 3C**). As observed for the cardiopharyngeal trajectory (**Figure 3-S1C**), pseudotime values were highly correlated with both real and inferred real times (**Figure 3-S1C**), indicating that our multiplexing approach faithfully recovered the transcriptome dynamics underlying developmental progression in the whole embryo. To facilitate exploration of these datasets, we created web-based interfaces using the ShinyApp package, and made them available at http://shiny.bio.nyu.edu/at145/whole_embryo and http://shiny.bio.nyu.edu/at145/tvc_atm_cells.

**Figure 3.**
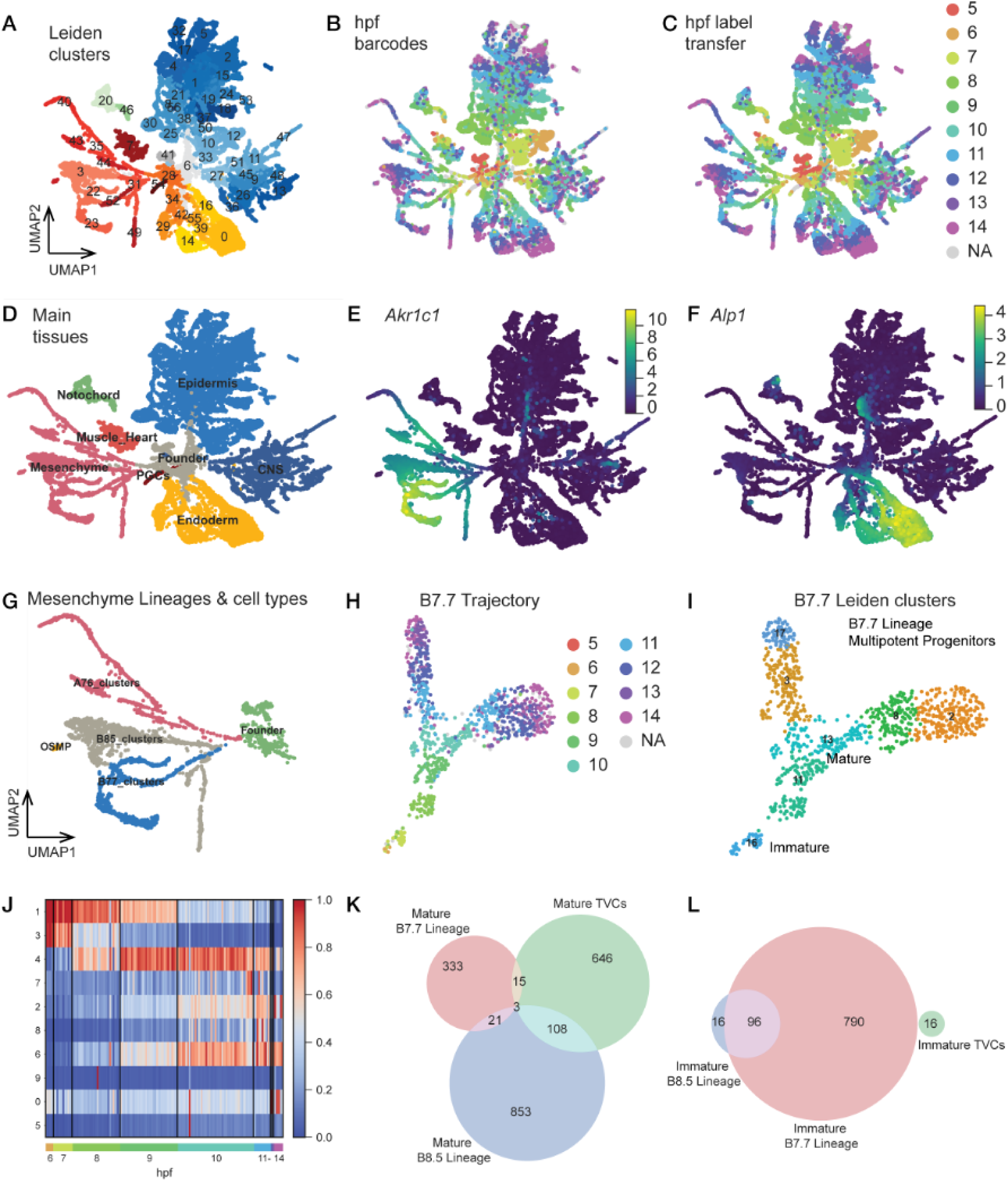
Whole embryo single-cell trajectories reveal maturation signatures in canalized endomesodermal lineages. **A.** Unifold manifold approximation and projection (UMAP) of the whole embryo scRNA-seq dataset colored and labeled by leiden clusters, with shades of blue for mesodermal epidermal and neuronal clusters, yellow for endodermal clusters, red for mesenchymal, germ line and muscle clusters and green for the notochord. The same umap is colored by developmental time obtained from the electroporated barcodes (**B**), or inferred through label transfer from closest neigbors (**C**). The same umap, colored by broad cell types, matching the color scheme detailed in a (**D**). **E-F.** Denoised expression of the pan-mesenchymal marker Akr1c1 (e; “KY.Chr1.2267”), and the pan-endoderm marker Alp1 (f; KY.Chr6.211”). **G.** Denoised Leiden clustering of mesenchymal tissues. **H,I.** Differentiation trajectory of the B7.7 mesenchymal lineage colored by inferred developmental time (H) and denoised leiden clustering (I). Note that cluster 16 corresponds to an immatue state and cluster 11 to a mature trancriptomic state. **J.** Differential expression of genes during differentiation of the B7.7 mesenchymal lineage. Gene clusters 1 and 4 correspond to the immature and mature states, respectively. **K,L.** Venn diagram showing the overlap of genes included in the mature (K) and immature gene clusters (L) of the mesenchymal B7.7 (red) and B8.5 (blue) lineages and the TVCs (green).

Remarkably, UMAP-based projection of the whole dataset illustrated cell fate diversification from early clusters located centrally toward diverging late cell identities located peripherally on the 2-dimension plot (**Figure 3A-C**). A cursory annotation using known markers identified the main tissue types throughout their transition from the gastrula to pre-hatching larval stage (**Figure 3D-F, Figure 3-S2**). Specifically, the notochord and muscle clusters, including a small number of B7.5/heart lineage cells, separated from the earliest time points onward, consistent with precocious cell fate specification, as early as the 64-cell stage, and post-mitotic differentiation during embryogenesis. Closer inspection of the notochord cluster suggested that its internal structure followed primarily the temporal sequence, without effectively distinguishing between the primary (A-line) and secondary (b-line) lineages. By contrast, and similar to a previous study^48^, the central nervous system clusters exhibited a complex structure that was only partially driven by temporal progression, and blurred the clonal distinction between A- and a-line derivatives. Likewise, the expansive and complex epidermal clusters separated by time points from 6 to 8 hpf, albeit displaying the premise of internal structure, whereas the 9 to 14 hpf cells diversified and clustered orthogonally to temporal progression, principally according to the dorsoventral and antero-posterior (e.g. tail-trunk) patterning, peppered with diverse epidermal sensory neurons. As for the central nervous system, initial clonal distinction between a- and b-line derived epidermal cells did not appear to be a main driver of transcriptome diversity, even at early time points.

By contrast with ectodermal clusters, endomesodermal lineages showed clearer signs of clonally-related developmental canalization over time (**Figure 3G-I**). For example, posterior B-line trunk endoderm emerged as early at 8 hpf, A-line endoderm and A7.6-line mesoderm (aka trunk lateral cells, TLCs) diverged transcriptionally as early as 6 hpf in gastrula embryos (**Figure 3**, **Figure 3-S2**). As for the muscle clusters, B-line mesenchymal lineages were already separated at the onset of the time series, and continued to diversify in progressively canalized manner toward defined late states.

Specifically, the B7.7 lineage (**Figures 3H, 3-S3**) appeared transcriptionally distinct from the B8.5 lineage as early as 6 hpf and further bifurcated into two sublineages, expectedly B8.13 and B8.14, around 10 hpf (mid-tailbud stage). Following an approach similar to that used for the cardiopharyngeal lineage (**Figure 2**), we identified three pre- and two post-bifurcation regulatory states for each branch. These transcriptional states coincided with 10 gene clusters including late, branch-specific clusters 2 and 6 (**Figure 3**, supplements). We further noted that cluster 4 and cluster 1 followed transient expression profiles, consistent with mature and immature states of the multi/bipotent B7.7 progenitors, respectively. Of note, as for the cardiopharyngeal progenitors, *Cdc25* [KY.Chr5.722] was part of the mature B7.7 genes (cluster 4) and transiently expressed shortly before the bifurcation (**Figure 3-S3**).

The B8.5 lineage followed a more complex trajectory profile, resulting in three transient and three distinct terminal transcriptional states of B8.5-lineage cells by late tailbud stages (**Figure 3I**, **Figure 3-S4**). The terminal transcriptional states (regulatory states 0, 15 and 25) were defined by *Irx6*, *Mist/Bhlha15* and *KY.Chr10.172*, respectively. Of note, B8.5-derived cluster 15 converges toward a transcriptional state that becomes indistinguishable from that of B7.7-lineage cells, including expression of *Mist/Bhlha15*. In total, genes expressed in this lineage could be grouped in 7 clusters, including immature cluster 4 and mature cluster 1 (**Figure 3-S3**).

As both mesenchymal and cardiopharyngeal lineages appear to transition through intermediate regulatory states before fate diversification, we sought to identify a gene expression signature associated with multipotent progenitor maturation in these distinct lineages. By intersecting candidate mature and immature markers of each lineage (**Figure 3K,L**), we identified only three candidate mature genes shared between all three lineages, namely (*Chdh-KY.Chr1.882*, *Prmt7-KY.Chr2.2197*, *Trp53i11-KY.Chr6.477*). By contrast, the B7.7 and B8.5 lineages shared 85.7% (97/112) of their candidate immature genes, consistent with their common origin as B-line mesenchyme progenitors. Thus, similar to the cardiopharyngeal progenitor trajectory, it appears that multipotent mesenchymal progenitors also progress along canalized trajectories that comprise transcriptional maturation prior to branching points and subsequent fate decisions.

### Mature state-specific determinants are required for heart vs. pharyngeal muscle fate choices

Having uncovered and begun to characterize transcriptome maturation in cardiopharyngeal and other multipotent progenitors, we sought to explore its biological significance. Our initial observations suggest that, although cardiopharyngeal progenitor cells could be forced to divide precociously, they were largely incapable of activating *Tbx1/10* until shortly before its normal timing in control embryos. We reasoned that, besides mitosis, transcriptome maturation might determine the competence of progenitor cells to activate *Tbx1/10* in addition to the known importance of Hand-r and FGF-MAPK signaling^37^.

To begin to test the hypothesis of a cell-autonomous competent state characterized by down-regulation of early markers and activation of late genes, we focused on candidate genes activated specifically toward the end of the cardiopharyngeal trajectory, in mature progenitor cells, which we hypothesized may be necessary for cardiopharyngeal fate decisions.

We selected 37 candidate genes, including those coding for such signaling molecules as the *trans*-membrane receptors *Adra2b* and *Fzd8*, or the Rho GTPase Activating Protein (RhoGAP) *Depdc1b*. We designed 111 single guide RNA (sgRNA)-expressing constructs for CRISPR/Cas9-mediated and B7.5 lineage-specific mutagenesis, using the *Mesp* enhancer to express SpCas9^49,50^. To assay phenotypes, we scored division orientation and asymmetry, and *Tbx1/10* expression at stage 25, after control cardiopharyngeal progenitors have normally divided asymmetrically along the medio-lateral axis, and the large lateral progenitors have activated *Tbx1/10* (**Figure 4A-B, Figure 4-S1**).

**Figure 4.**
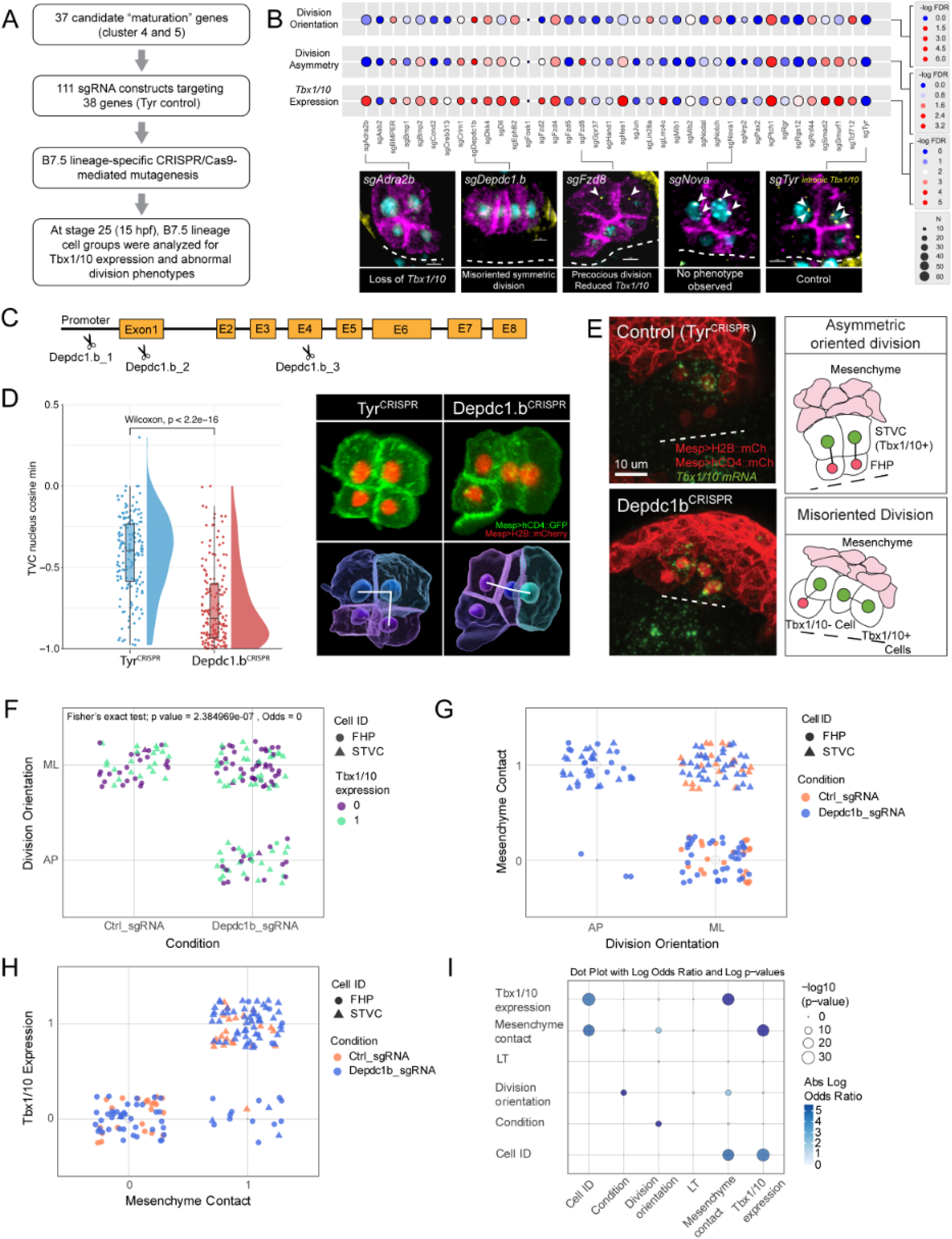
Maturation state-specific determinants are required for pharyngeal vs heart muscle fate choices. **A.** Flowchart of the CRISPR screen library generation and embryo collection. **B**. Phenotypes associated with CRISPR targeting of indicated loci. −log of the false discovery rate (FDR) is shown with a cutoff value of 0.05. Micrographs show examples of phenotypes produced. Membranes are marked with lineage specific Mesp>hCD4::mCh (magenta), nuclei with Mesp>LacZ (cyan), and a DIG-labeled intronic RNA probe against Tbx1/10 transcripts (yellow). **C**. Map of the Depdc1.b genomic locus with indicated guide RNA targets. **D**. Range of phenotypes produced by CRISPR disruption of the Depdc1.b locus. Graph shows the change in asymmetric division as ratios of FHP to STVC volumes and change in division orientation as the cosine of the angle between cell pairs. Micrographs show rendered volumes of cells color coded by relative volume and division orientation indicated by white lines. **E**. Micrographs showing FISH targeting the *Tbx1/10* transcripts (green) in the FHP and STVCs. Mesenchymal tissue is marked with tissue specific Twist>hCD4::mCherry. STVC and FHP membranes and nuclei are marked with Mesp:hCD4::mCh and Mesp>H2B::mCh respectively. Dashed lines mark the embryonic midline. Diagrams to the right highlight the contact between Tbx1/10 expressing cells and the mesenchyme. **F,G,H.** Correlation of indicated parameters. F. CRISPR targets and division orientation, G. Division orientation and mesenchymal contact, and H. Tbx1/10 expression. **I**. Log odds ratios and significant associations of characteristics and phenotypes between control and Depdc1b CRISPR conditions.

Treating division orientation and asymmetry, and *Tbx1/10* expression as categorical variables, we applied Fisher’s exact tests and calculated false discovery rates (FDR) to identify the diverse combinations of selected phenotypes induced by defined perturbations. For example, CRISPR/Cas9 mutagenesis of *Adra2b* significantly inhibited *Tbx1/10* activation without affecting the cell division patterns, while targeting Frizzled-coding genes variably altered cell divisions and/or *Tbx1/10* expression. In total, we identified 25 out of 37 candidate “mature genes” causing a *Tbx1/10* expression and/or cell division phenotype following CRISPR/Cas9-mediated loss-of-function (**Figure 4B**). The variable combinations of phenotypes observed hint at the partial independence of the subcellular phenomena, and thus the modulatory of biomolecular networks governing *Tbx1/10* expression and division orientation and asymmetry. On the other hand, with the exception of *Lin28*, all conditions altering division orientation and/or asymmetry also impacted *Tbx1/10* expression. This suggested that, beyond the system’s modularity, the geometry of cardiopharyngeal progenitor divisions influence subsequent *Tbx1/10* expression patterns.

Focusing on the DEP domain-containing RhoGAP-encoding *Depdc1b*, fluorescent *in situ* hybridization validated its expression in maturing progenitor cells, starting approximately at the G1-S transition, and confirming that *Depdc1b* activation precedes oriented and asymmetric divisions in late tailbud embryos (**Figure 2-S2**, see also **Figure 5A**). To further characterize the *Depdc1b^CRISPR^* division phenotype, we imaged, segmented and quantified defined morphometric parameters in both control, *Tyrosinase^CRISPR^*, and *Depdc1b^CRISPR^* cardiopharyngeal lineage cells (**Figure 4C,D**), as part of systematic high-content image-based CRISPR screen (Failla, Wiechecki et al., *in preparation*). Specifically, volumetric ratios and angles between axes formed by cell triplets, which report on asymmetry and orientation of cell divisions, respectively, indicated that the control *Tyrosinase^CRISPR^*and *Depdc1b^CRISPR^* embryos occupy distinct territories in this simple 2-dimension phenotypic space. This suggested that Depdc1b function is required for both the asymmetry and orientation of cardiopharyngeal progenitor divisions. This role is reminiscent of the function of *LET-99*, the *Depdc1b* homolog in *Cænorhabditis elegans*, which functions to position and orient the mitotic spindle during the first embryonic cleavage^51,52^.

**Figure 5.**
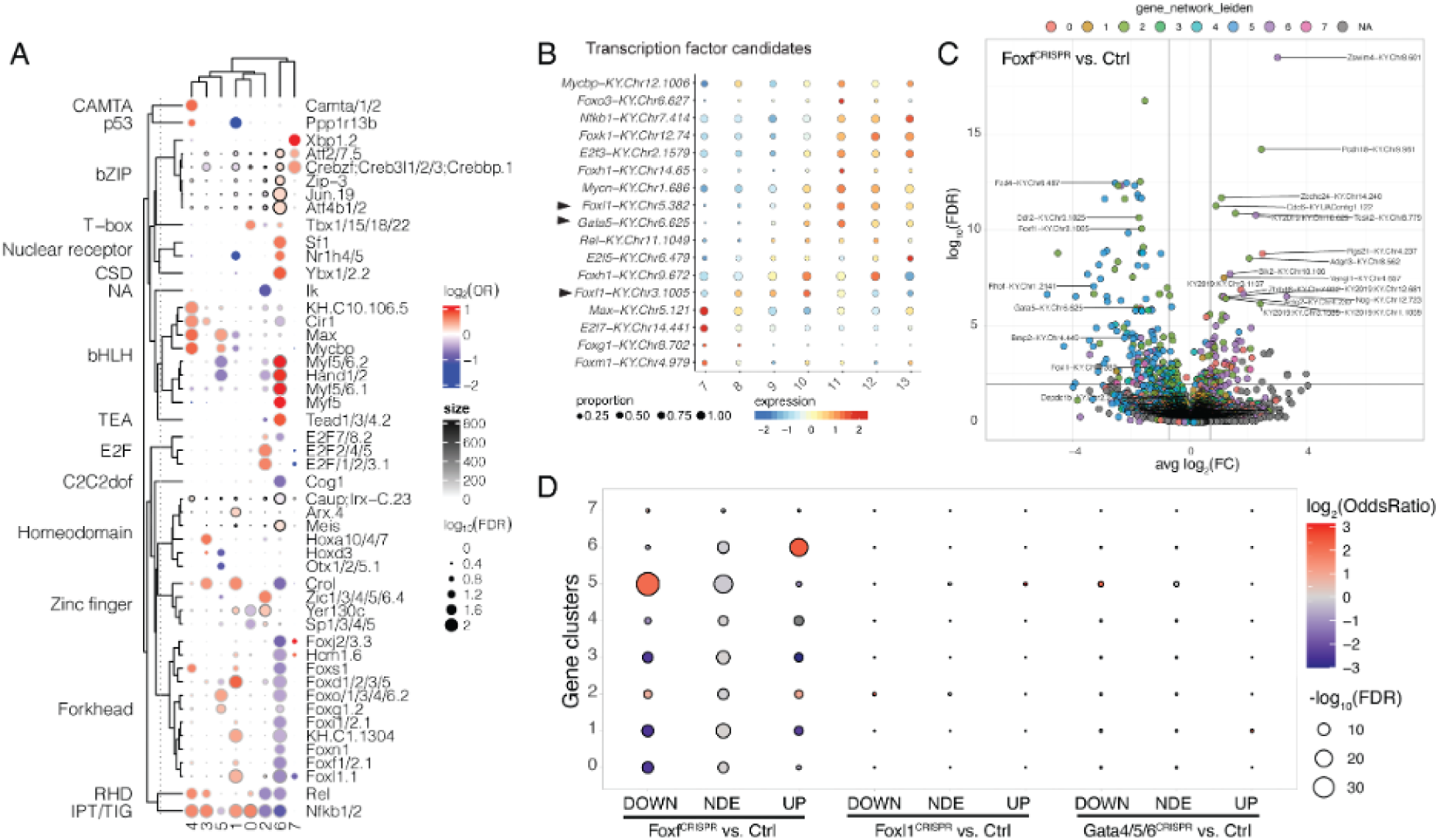
Candidate transcription factors involved in maturation gene regulation. **Candidate transcription factors involved in the regulation of maturation genes** **A.** Cluster-specific transcription factor motif enrichment. Dot color gives the log2 odds ratio of occurrence of a motif in a motif set to the occurrence of a motif in the background. Dot size gives the p-value of the hypergeometric test. Outline color gives occurrences of a motif in the motif set. **B.** Expression dynamics of candidate transcription factor expression mapped on real-time barcodes. Arrowheads indicate CRISPR targets used in C and D. **C.** Volcano plot showing differentially expressed genes between control (Ctrl) and Foxf CRISPR conditions. Log fold change is shown on the x-axis. Color code represents gene clusters as shown in figure 2i. **D.** Differential expression of genes grouped by gene cluster between control (Ctrl) and indicated CRISPR conditions.

Remarkably, even though Depdc1b controls the orientation and asymmetry of cardiopharyngeal progenitor divisions, its disruption caused gene expression and fate decision phenotypes following abnormal divisions (**Figure 4E-I**. Specifically, *Dedpc1b^CRISPR^* caused a significant fraction of sister cell pairs to align along the antero-posterior axis, by contrast with the medio-lateral alignment typically observed in control embryos (**Figure 4F**). Sister cells misalignment significantly correlated with increased contact with the lateral *Twist>hCD4::mCherry*+ mesenchymal lineages (**Figure 4E,G**), which in turn correlated significantly with *Tbx1/10* expression (**Figure 4E,H**). Despite the variability and incomplete penetrance of the *Depdc1b^CRISPR^* phenotype, these analyses thus indicate that Depdc1b control the orientation of cardiopharyngeal progenitor divisions, which in turn determines differential contact with the lateral mesenchyme and polarized activation of *Tbx1/10* specifically in the lateral second-generation cardiopharyngeal progenitors.

To further explore the role of Depdc1b in cardiopharyngeal progenitor divisions we re-evaluated the Cdc25^OE^ phenotype, which was characterized by precocious but also misoriented and/or symmetrical divisions, in a manner reminiscent of the *Depdc1b^CRISPR^* phenotype (**Figure 1-S2**). Since *Depdc1b* expression starts at the G1-S transition and peaks during G2 in the mature state, we reasoned that Cdc25^OE^ may force cells to divide before Depdc1b reaches a functional level, thus impairing proper oriented and asymmetric division. Consistent with this hypothesis, Cdc25^OE^ did not impair the onset of *Depdc1b* expression in early S-phase, but the shortened G2 phase, causing cells to divide with lower *Dedpc1b* expression level, consistent with a contribution to the Cdc25^OE^ cell division orientation and asymmetry phenotypes (**Figure 4-S1**).

Taken together, these results indicate that *Depdc1b* upregulation endows mature cardiopharyngeal progenitors with the ability to divide in an asymmetric and oriented manner, allowing them to produce distinct large *Tbx1/10*+ second cardiopharyngeal progenitors, laterally, and small *Tbx1/10-* first heart precursors, medially. We propose that transcriptome maturation of progenitors foster the cellular competence to generate both cardiac and pharyngeal muscle lineages, a hallmark of multipotent cardiopharyngeal progenitor cells.

### Foxf and Gata-driven feedforward sub-circuits promote multipotent progenitor maturation

The above results indicated that the transcriptome of multipotent progenitors changes as they migrate and progress through the cell cycle. This molecular maturation fosters the competence to undergo oriented and asymmetric divisions that condition the production of distinct cardiac and pharyngeal muscle lineages. Considering the importance of multipotent progenitor maturation for subsequent fate decisions, we sought to explore the mechanisms governing transcriptome dynamics during maturation.

We previously described the changes in the immediate environment of multipotent progenitors, which are born laterally, in the vicinity of the mesenchyme and notochord, before migrating ventro-medially in between the epidermis and trunk endoderm^40,53^. It is likely that distinct surrounding tissues exert variable influence on migrating progenitors. Nevertheless, here we focused on possible cell-autonomous drivers of transcriptome dynamics.

Leveraging our previously published B7.5-lineage-specific chromatin accessibility data^54^, we sought to identify sequence motifs enriched in accessible regions surrounding genes clustered by pseudotemporal profiles along the cardiopharyngeal trajectory (**Figures 5A, 2H**). Consistent with their roles in tail muscle specification, this analysis identified motifs for the myogenic transcription factor Myf5/Myod in cluster 6, which corresponds to genes maintained specifically in the anterior tail muscle cells (**Figure 2-S2D**). By contrast, Forkhead/Fox family motifs were depleted in the accessible regions associated with this gene cluster, also consistent with the previously recognized role for Fox family factors in cardiopharyngeal fate choices^41,54,55^.

As candidate cell-autonomous regulatory mechanisms, we considered classic gene regulatory network motifs such as feedforward circuits^56^. Previous work identified *Foxf* as one of the earliest developmental control genes activated in cardiopharyngeal progenitors, in response to FGF-MAPK signaling^41,46^. Foxf controls subsequent activation of the conserved cardiac transcription factors coding genes *Gaca4/5/6*, *Nk4/Nkx2.5* and *Hand*^41,46,55^, at least in part by promoting chromatin accessibility at cardiopharyngeal enhancers, and subsequent transcriptional activation^54^. Transcription factors of the GATA4/5/6 family also play essential and conserved regulatory roles in cardiac vs. pharyngeal lineage specification^8–10^. To assay the roles of candidate TFs during maturation of cardiopharyngeal progenitors, we performed CRISPR/Cas9-mediated mutagenesis of Foxf and Gata4/5/6 along with a forkhead family transcription factor Foxl1, among other candidate TFs dynamically expressed in the cardiopharyngeal trajectory (**Figure 5B**). We profiled the transcriptomes of FACS-purified cells at stage 23/24 (∼12 hpf at 18°C), when control cells reach the mature state, using transgenic barcode-mediated multiplexed scRNA-seq. We pooled four barcoded samples corresponding to *Foxf*, *Gaca4/5/6*, *Foxl1*-specific and control CRISPR/Cas9 reagents into a single library for scRNA-seq, followed by *in silico* demultiplexing.

Building on highly correlated barcode pairs (**Figure 5-S1A**), we confidently assigned 112, 126, 92 and 193 single cell transcriptomes to either *Foxf*, *Gaca4/5/6*, *Foxl1* loss-of-function or control conditions, respectively (**Figure 5-S1B**). Among those, clustering, UMAP projection and marker gene expression identified 223 cardiopharyngeal progenitors and 326 anterior tail muscle, indicating that reporter-driven barcodes are effective to identify B7.5-lineage cells with >95% certainty at early stages (**Figure 5-S1C-E**). *In silico* demultiplexing of pooled scRNA-seq then allowed us to evaluate the cardiopharyngeal progenitor-specific effects of individual perturbations independently of cell sorting (**Figure 5-S1F,G**). *Foxf*, *Gaca4/5/6*, and *Foxl1* were among the most significantly down-regulated genes in their corresponding CRISPR/Cas9 conditions (**Figure 5C**, **Figure 5-S2A,B**), indicating effective perturbations. Both *Gaca4/5/6* and *Foxl1* were also down-regulated in *Foxf^CRISPR^* cells, consistent with previous observations and the established role for Foxf in driving chromatin accessibility and gene activation in the cardiopharyngeal lineage (**Figure 5C, Figure 5-S2**). Applying permissive statistical cutoffs (FDR < 0.05 and |Log_2_(FC)| > 0.5), we identified 302 candidate differentially expressed genes in *Foxf^CRISPR^* compared to control cells, including 192 down- and 110 upregulated transcript. By contrast, *Gaca4/5/6^CRISPR^* and *Foxl1^CRISPR^* yielded more modest numbers, with 9 and 1 gene significantly). Despite a modest genome-wide impact, the top candidate *Gaca4/5/6* target was *Depdc1b*, the mature-state marker required for oriented asymmetric division (**Figure 5-S2A**, see also **Figure 6** below).

**Figure 6.**
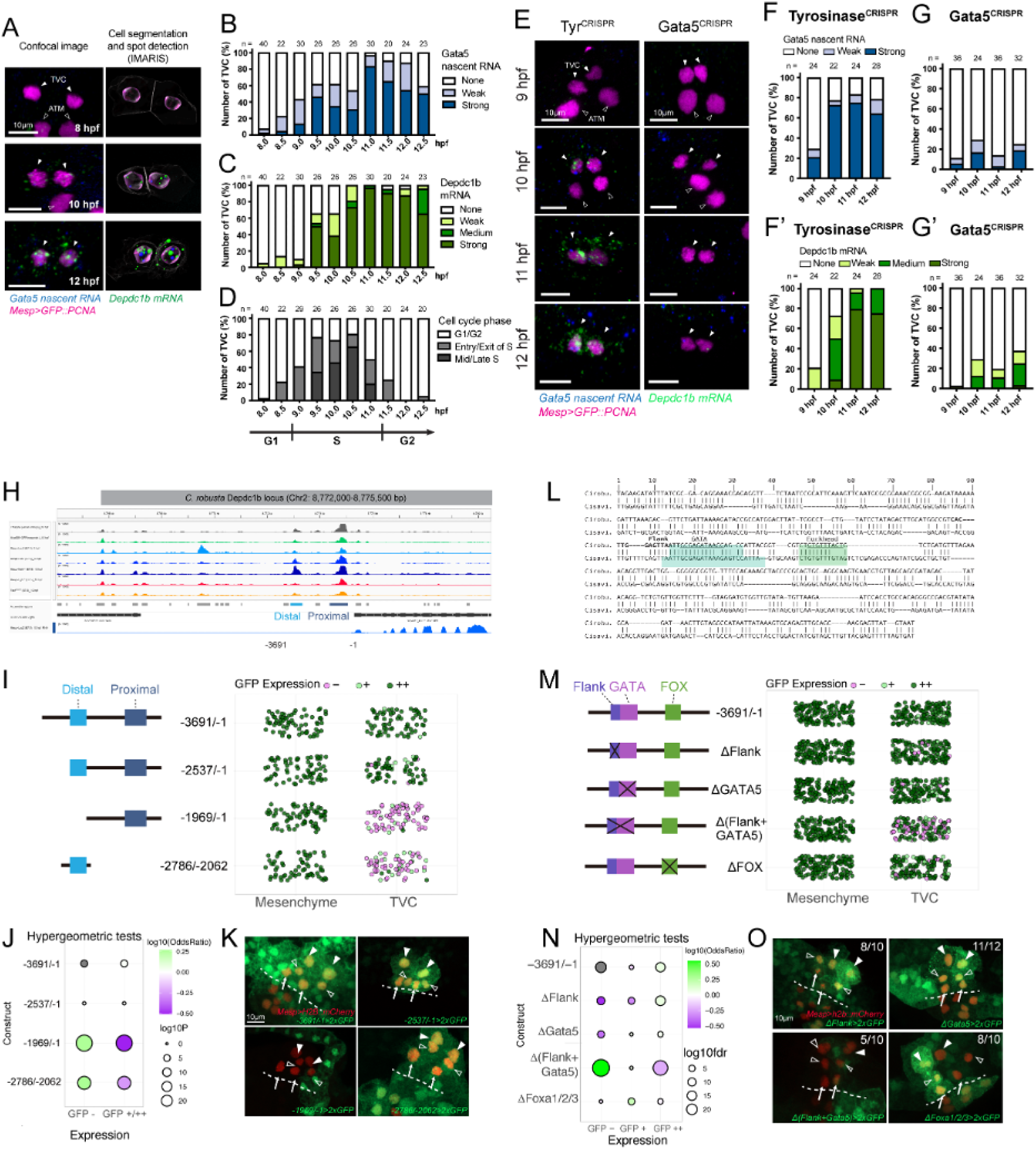
Gata4/5/6 regulates Depdc1b expression. **A-C.** Developmental dynamics of Gata4/5/6 gene expression. a. Representative confocal images showing double FISH detection of Gata4/5/6 and Depdc1b expression. Segmented cells, nuclei and transcripts shown on the right. Magenta: nuclei (GFP::PCNA); Blue: Gata5 nascent RNA; Green: Depdc1b mRNA. Scale bar = 5mm; Arrowhead: TVC; Open arrowhead: ATM. B-C. Quantification of Gata4/5/6 (B) and Depdc1b (C) gene expression spanning TVC migration but prior to TVC division. **D.** Cell cycle phases as determined by the GFP::PCNA cell cycle reporter. **E.** Confocal images of double FISH targeting Gata4/5/6 and Depdc1b under control (TyrCRISPR) and Gata4/5/6CRISPR knockouts. Scale bar = 10 um Arrowhead: TVC; Open arrowhead: ATM. **F-F’** and **G-G’.** Quantification of Gata4/5/6 (F-F’) and Depdc1 (G-G’) expression in control (TyrCRISPR) and Gata4/5/6CRISPR knockout condition. **H.** Accessibility of the Depdc1b locus during development showing consensus accessible regions and highlighting distal and proximal regulatory regions upstream of the Depdc1b start site. **I.** Systematic deletion of distal and proximal regulatory regions upstream of a 2xGFP reporter. Diagram shows control and deletion constructs. Dot plot on the right shows level of GFP expression detected in either the Mesenchyme or the TVCs upon disruption of regulatory regions, separately or in combination. Jitter is added to the graph for ease of visualizing expression changes. **J.** Hypergeometric tests of GFP expression based on regulatory region perturbation. Color scale indicates log10 odds ratio and size indicates log10 p-value. **K.** Micrographs of GFP expression as driven by constructs containing both proximal and distal regulatory regions or lacking one or both regulatory regions. **L.** Regions of the Depdc1b regulatory region showing conservation of the Flank region, GATA binding side, and Forkhead binding site between *Ciona robusta* and *Ciona savignyi*. **M.** Constructs containing the Flank, GATA, and FOX binding sites driving 2xGFP expression and subsequent analysis of binding site requirements for the tissue specific expression of GFP. Dot plot on the right shows level of GFP expression detected in either the Mesenchyme or the TVCs upon disruption of regulatory regions, separately or in combination. Jitter is added to the graph for ease of visualizing expression changes. **N.** Hypergeometric tests of GFP expression based on regulatory region perturbation. Color scale indicates log10 odds ratio and size indicates log10 p-value. **O.** Micrographs of GFP expression as driven by constructs containing both proximal and distal regulatory regions or lacking one or both regulatory regions. For both K and O, Arrows – First Heart Precursors (FHPs), open arrow heads – Second Heart Precursors (SHPs), solid arrow heads Atrial Siphon Muscle Founder cells (ASMF).

Leveraging its more widespread impact on the cardiopharyngeal transcriptome (and chromatin accessibility^54^), we focused on the *Foxf^CRISPR^*condition to evaluate its role(s) in transcriptome maturation. First, we verified that the multiplexed scRNA-seq approach correlated with previous profiling of the same *Foxf^CRISPR^* perturbation by bulk RNA-seq^54^ (Pearson’s *𝛠* = 0.91, *p* < 2.2 x 10^−16^;). Next, we used the *Foxf^CRISPR^* vs. control volcano plot to visualize the individual responses of genes grouped by pseudotemporal clusters along the cardiopharyngeal progenitor trajectory (**Figure 5C**). This suggested that genes in cluster 5, which comprises mature-state-specific markers (**Figure 2I**), are mostly down-regulated, while the intermediate gene cluster (cluster 2; **Figure 2I**) splits between up- and down-regulated genes. On the other hand, cluster 6, which comprises anterior tail muscle markers down-regulated in cardiopharyngeal progenitors, comprised genes that were upregulated (**Figure 2-S2D**). Hypergeometric tests showed that clusters 2 and 5 were significantly over-represented among down-regulated genes, while clusters 2 and 6 were over-represented among up-regulated genes (**Figure 2D**). Taken together, these observations indicate that the early cardiopharyngeal determinant Foxf, promotes multipotent progenitor maturation both by activating intermediate and mature-state markers and by down-regulating other intermediate markers as well as the alternative anterior tail muscle program (see discussion).

As indicated above, feed-forward circuits are classic and wide-spread gene regulatory network motifs accounting for the deployment of lineage specific transcriptional programs in development^57^. The above and previous analyses placed Foxf atop the cardiopharyngeal regulatory hierarchy in Ciona, including *Gaca4/5/6* as one of its notable target genes^41,46,54,55^ (**Figure 5C**).

Intriguingly, the mature gene *Depdc1b*, which is necessary for proper oriented and asymmetric division of cardiophayrngeal progenitors (**Figure 6A-D**), was also the top down-regulated gene in *Gaca4/5/6^CRISPR^* cells, prompting us to further explore the regulatory relationship between the two genes. First, double FISH assays using embryos expressing the S-phase *Mesp>Pcna::GFP* indicated that *Gaca4/5/6* expression precedes that of *Depdc1*, which started at the G1-S transition (**Figure 6A-D**). Double FISH assays following CRISPR/Cas9-mediated loss of *Gaca4/5/6* function corroborated the scRNA-seq data, whereby both *Gaca4/5/6* and *Depdc1b* were strongly down-regulated in *Gaca4/5/6^CRISPR^* embryos. These data indicated that Gata4/5/6 funcion is necessary for *Depdc1b* expression, which starts at the G1-S transition.

Leveraging our previous cardiopharyngeal lineage-specific chromatin accessibility data^54^, we identified 2 main accessible regions located approximately 2.5 and 0.5 kbp upstream of the *Depdc1b* start site (**Figure 6H**). Notably, the region located at ∼2.5 kb upstream displayed the typical ATAC-seq pattern of Foxf-dependent cardiopharyngeal progenitor-specific accessibility: the signal was higher in B7.5 lineage cells than whole embryos or mesenchyme lineage cells at 10 hpf, it increased between 6 and 10 hpf, and it was down-regulated in conditions that inhibit cardiopharyngeal progenitor induction or Foxf function. To test the importance of these elements for *Depdc1b* regulation, we built a reporter construct by cloning a 3,961 bp fragment upstream of a 2xGFP reporter, which we electroporated into fertilized eggs alongside a *Mesp>H2B::mCherry* lineage marker and assayed GFP expression at stage 25, shortly after the division of cardiopharyngeal progenitors (**Figure 6I,J**). The original −3,961/-1 construct, and its trimmed −2,537/-1 version, showed similar GFP expression in the cardiopharyngeal lineage with 100 ± 0 % (n = 70; ± S.E.) and 92.8 ± 13.1% (n = 69; ± S.E.) of the embryos, respectively (Hypergeometric test p = 0.028; **Figure 6J-K**,). By contrast, deletion of either the distal or proximal elements significantly reduced the proportions of embryos with detectable expression in the cardiopharyngeal lineage to 16.9 ± 4.4 % (n = 71; ± S.E.; p = 2.6 x 10^−21^) and 35.9 ± 6.0 % (n = 64; ± S.E.; p = 1.3. x 10^−12^), respectively (**Figure 6I-L**, Table S). These results indicate that both distal and proximal elements are necessary for cardiopharyngeal expression of the *Depdc1b* reporter construct.

Next, we focused on the distal element, which showed cardiopharyngeal-specific and Foxf-dependent accessibility. This element proved to be highly conserved between *Ciona robusca* and its sibling species *Ciona savignyi*, with short blocks of identical sequences that typically correspond to putative transcription factor binding sites (**Figure 6M**). Joint motif search using both sequences and the Cis-BP database and software^58^ identified several high-scoring putative binding sites for such cardiopharyngeal regulators as ETS, bHLH, Homeodomain, FOX and GATA family factors (**Figure 6-S1**). Focusing on the latter, we introduced microdeletions of conserved sequences corresponding to putative GATA and FOX binding sites, and a conserved region flanking the GATA motif in the original −3961/-1 construct, and assayed reporter gene expression (**Figure 6N-Q**). Of these, the joint “Flank + GATA” deletion caused a drop of cardiopharyngeal expression in electroporated embryos to 59.0 ± 4.2 % (n = 139; ± S.E.) from 96.6 ± 1.5 % (n = 146; ± S.E.) and 95.3 ± 1.5% (n = 148; ± S.E.) with constructs lacked either the “Flank” or GATA motif, respectively (**Figure 6**;). Taken together, these data suggest that Gata4/5/6 activate *Depdc1b* expression in cardiopharyngeal progenitors, at least in part by directly binding to the evolutionary “Flank+GATA” motif located in a lineage-specifically accessible distal upstream element.

### Cell cycle progression promotes *Depdc1b* expression

Finally, several of the above observations led us to reason that cell cycle progression through interphase could promote progenitor maturation. Specifically, the mature state emerges during the late G2 phase, and marked dynamics in transcriptome composition coincide with phase transitions, especially G1-S, which also correlates with the onset of both collective migration and *Depdc1b* expression (**Figures 1, 2**). Focusing on the impact of the G1-S transition, we inhibited it by overexpressing Cdnk1b (Cdkn1b^OE^) in the cardiopharyngeal progenitors using the minimal *Foxf* enhancer as previously described^37^ (**Figure 7A**). We assayed the Cdkn1b^OE^ phenotypes using the S-phase reporter Mesp>GFP::PCNA and double FISH to detect *Gaca4/5/6* and *Depdc1b* transcripts (**Figure 7B**). As expected, Cdkn1b^OE^ significantly reduced the proportions of cardiopharyngeal progenitors entering S-phase during the 9 to 12 hpf window observed in controls (**Figure 7C,D**). This perturbation did not alter the onset *Gaca4/5/6* expression, which remained detectable as early as 9 hpf, even though it did lower its highest level (**Figure 7B,C’D’**). On the other hand, Cdkn1b^OE^ markedly inhibited *Depdc1b* activation, as well as its accumulation toward the end of interphase (**Figure 7B,C”,D”**). Together, these observations provide direct experimental support to the notion that progression through G1-S promotes transcriptome maturation in cardiopharyngeal progenitors as exemplified by the impact of Cdkn1b^OE^ on the activation of *Depdc1b*, a mature state marker required for proper oriented and asymmetric division of multipotent progenitors.

**Figure 7.**
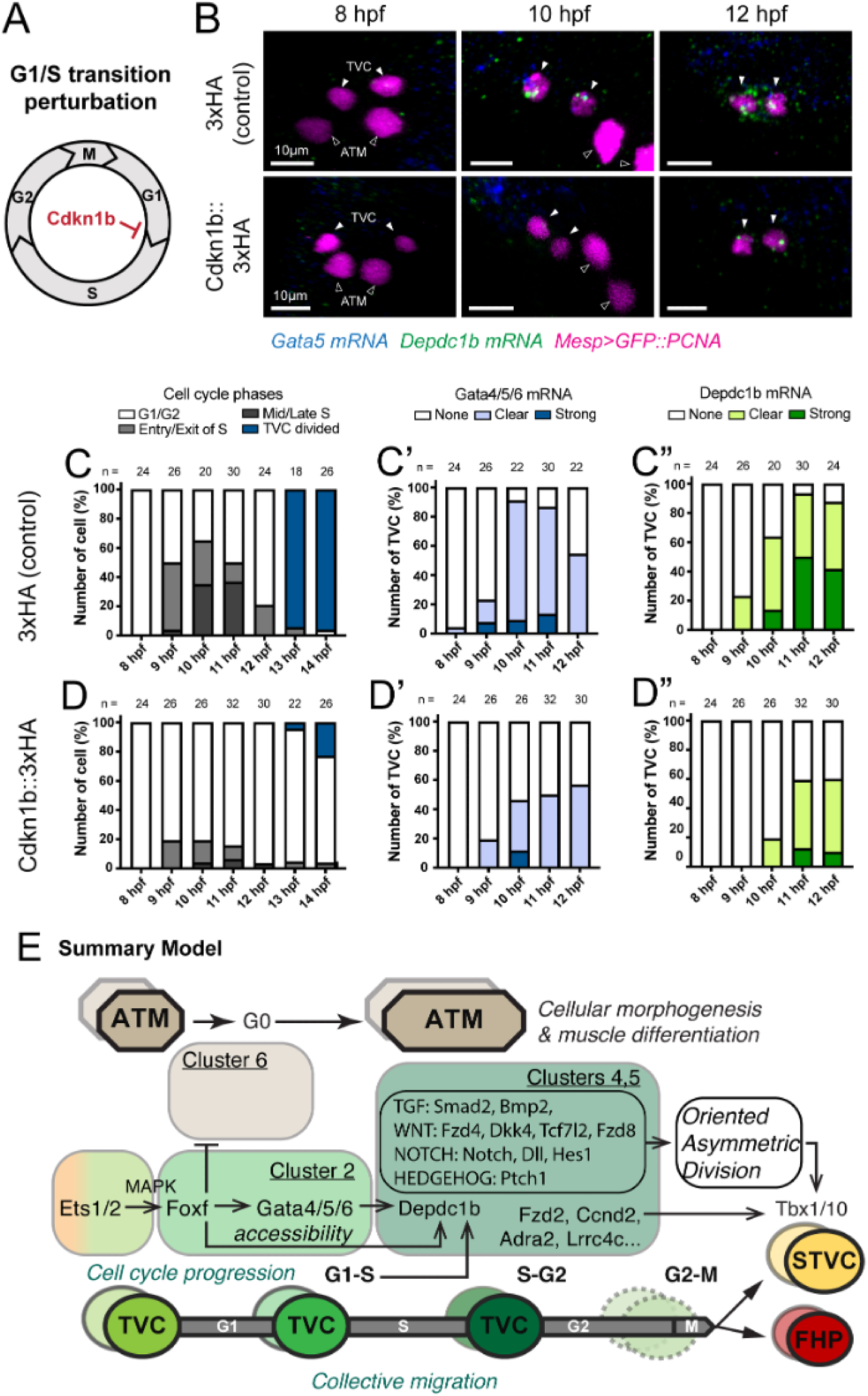
Cell cycle progression drives feed-forward sub-circuits and promotes multipotent progenitor maturation. **A.** Schematic of cell cycle stages and genetic perturbation of G1/S transition. **B.** Representative confocal images of Gata4/5/6 and Depdc1b expression at 8, 10, and 12 hpf under control and Cdkn1b::3xHA^OE^ conditions. Magenta: nuclei (GFP::PCNA); Blue: Gata5 mRNA; Green: Depdc1b mRNA. Scale bar = 10mm; Arrowhead: TVC; Open arrowhead: ATM. **C-D.** Dynamic expression of Gata4/5/6, Depdc1b expression, and S phase progression in the control and Cdkn1b-overexpressing TVCs. **E.** Working model of the feed-forward sub-circuits driven by cell cycle progression.

## Discussion

As development proceeds, cell identities are canalized through progressive fate restriction of multipotent progenitors. Focusing on multipotent cardiopharyngeal progenitors in the tunicate *Ciona*, we explored the acquisition of multilineage competence and the coupling of progressive fate specification with cell cycle progression. Combining defined perturbations of cell cycle progression with gene expression analysis, we first provided evidence that mitosis is required but not suficient for *Tbx1/10* activation, an essential step in the heart vs. pharyngeal muscle fate choice^35–37,39^. Using sample multiplexing and scRNA-seq to profile a time-series of FAC-sorted cardiopharyngeal progenitors, we uncovered transcriptome dynamics characterized by transitions through regulatory states, which we termed “maturation”. We identified gene clusters representing defined (pseudo)temporal profiles in the maturation trajectory. The S-phase marker PCNA::GFP revealed a correspondence between the G1-S and S-G2 transitions and the maturation of multipotent progenitors, which reach a mature state characterized by the down-regulation of early and intermediate markers and peak expression of “late/mature genes” in late G2. Using systematic loss-of-function by CRISPR/Cas9-mediated mutagenesis, we showed that most “mature genes” are necessary for proper oriented and asymmetric division of multipotent progenitors, and subsequent polarized activation of *Tbx1/10*. Focusing on the RhoGAP-coding gene *Depdc1b*, which is activated at the G1-S transition, we uncovered a role in the orientation and asymmetry of cell division, which impacts the ability of multipotent progenitors to generate two distinct, *Tbx1/10*-positive and negative, daughter cells along the medio-lateral axis. These results supports the notion that multipotent cardiopharyngeal progenitors need to “mature” to acquire the competence to divide in an oriented and asymmetric fashion, and produce both a large lateral *Tbx1/10*+ second cardiopharyngeal progenitor and a small median first heart precursor. Finally, we present evidence that transcriptome maturation relies, at least, on classic gene regulatory network motifs such as feed-forward circuits, as well as coupling with cell cycle progression, especially the G1-S transition.

Classic concepts in developmental biology, dating from homo- and heterotopic and -chronic graftings experiments by gifted embryologists, suggested that progenitor cells transition through specification and commitment to restricted identities, while the notion of maturation has been considered extensively for differentiated cells, such as cardiomyocytes, which are contractile but not fully functional until they reach a mature state. Here, we adopted a general definition of the concept of maturation, considering a biological entity that persists through time and progressively acquires its full functionality through underlying changes. From that stand-point, since multipotent cardiopharyngeal progenitors transition through successive regulatory states as they progress through interphase, and ultimately acquire the competence to divide in an oriented and asymmetric manner and produce distinct first heart vs. *Tbx1/10*+ second craniopharyngeal progenitors, we argue their transcriptome maturation fosters multilineage competence.

In an attempt to generalize the notion of multipotent progenitor maturation, we leveraged our transgene-base barcoding strategy for multiplexed scRNA-seq, and profiled whole embryos from gastrula to pre-hatching larval stages in one experiment. With ∼22,000 single cell transcriptomes, we recovered the same main lineages and identities as a previous more extensive study, but with reduced batch effects. Our barcoded time stamped unequivocally identified developmental trajectories that followed temporal sequences. For instance, the transcriptional signatures of early lineages appeared overridden by patterning signatures in the ectoderm-derived epidermis and nervous system. By contrast, endomesodermal lineage displayed more conspicuous signs of developmental canalization, due to the persistence of transcriptional signatures of their clonal origins. Through subsetting and reclustering, we characterized specific trajectories, which we interpreted as multipotent progenitors “maturing” before splitting into distinct identities, and identified corresponding candidate mature gene clusters.

Extending beyond ascidian embryogenesis, whole animal profiling of developmental trajectories during vertebrate embryogenesis has repeatedly uncovered similar patterns of multipotent progenitors seemingly “maturing” along linear trajectories before producing distinct cell identities, presumably following cell divisions^59–61^. As we observed by comparing distinct trajectories within the ascidian embryo, gene expression signatures of maturing multipotent progenitors tend to be lineage and/or fate-specific. For instance, multilineage transcriptional priming is a hallmark of multipotent progenitor transcriptomes, and we observed that primed heart vs. pharyngeal muscle-specific genes are enriched in mature gene clusters. However, multilineage transcriptional priming started prior to the mature state as shown by another enrichment of primed genes in the intermediate gene cluster (cluster 2 in Figure 2), indicating that priming is completed through maturation, consistent with the notion that maturation corresponds the acquisition of full functionality. This indicates that multilineage transcriptional priming is one of the molecular system’s features that is acquired at least in part through transcriptome maturation.

Consistent with marked lineage-specific components to transcriptome maturation, classic gene regulatory network motifs such as feed-forward circuits contribute to progenitor maturation. Specifically, the extensive roles of key cardiopharyngeal determinants Foxf and Gata4/5/6 corroborates previous studies, as does the observed sequence of gene activation for primed cardiac determinants *Gaca4/5/6*, *Nk4/Nkx2.5*, and *Hand1/2*, which also depends on chromatin priming through enhancer opening by Foxf^54^. There is extensive evidence for such “regulatory competence”, whereby chromatin and transcriptional mechanisms determine multilineage competence, in a variety of developmental systems. Here, we showed that Gata4/5/6 also regulates the timed expression of the “mature gene” *Depdc1b*, which encodes a Rho GAP necessary for proper asymmetric and oriented division. Careful analysis of the CRISPR/Cas9-induced phenotypes suggested that Depdc1b controls primarily division orientation and asymmetry, which in turn impacts contact with the surrounding mesenchyme, and the pattern of *Tbx1/10* expression. This interpretation is consistent with the known roles of the Depdc1b homolog in *C. elegans*, LET-99, which controls the position and orientation of the mitotic spindle during the first embryonic division^51,52^. Taken together, these results indicate that transcriptome maturation also endows multipotent progenitors with “behavioral competence”, i.e. the cytoskeletal/cellular ability to divide in a manner compatible with the emergence of two distinct cell identities marked by differential *Tbx1/10* expression. Future work will uncover the role(s) of the lateral mesenchyme as a cardiopharyngeal niche (Christiaen lab., unpublished observations).

Our results indicate that cell cycle progression influences cardiopharyngeal fate specification in at least two distinguishable ways: *Tbx1/10* activation shortly follows mitosis, which is necessary, but not sufficient as multipotent progenitors themselves need to progress through G1, S and G2 to reach a mature state competent to express *Tbx1/10*. As reported above, the competence for polarized *Tbx1/10* activation requires oriented asymmetric division, but certain “mature genes”, such as the GPCR-coding genes *Adra2b* and *Lrrc4c* appeared to impact *Tbx1/10* without altering the division pattern. Given the short estimated delay between mitosis and *Tbx1/10* activation (<30 minutes), we anticipate gene activation to occur in G1. This is reminiscent of a CDK-independent and nuclear role for late G1 cyclin Ds in activating *TBX1* in human pluripotent stem cells^62^, and of the general propensity of the G1 phase to promote fate choices, for example during stem cell specification^14,63^. On the other hand, a high-throughput RNAi screen identified G2 phase regulators as promoters of the undifferentiated state in mammalian stem cells^64^. Here we propose that the transcriptome changes that unfold in G2 promote maturation of the multipotent progenitors, and their competence to produce distinct cardiopharyngeal cell identities. From this stand-point, we propose to reconcile the two views, since maturation completed the multilineage priming of progenitors, and argue that progression through G2 and maturation poise the cells to choose a cardiopharyngeal identity in subsequent divisions while preserving multipotency, and the competence to choose either fate. Finally, the tight coupling between cell cycle progression and fateful gene expression changes observed in ascidian lineages might represent an extreme case, in keeping with their highly stereotyped embryogenesis^65,66^. Recent work in *Drosophila* has also begun to uncover the relationship between mesodermal fate decisions, cell cycle progression and collective migration^67^. By contrast, the more relaxed coupling observed in vertebrates, together with signs of progenitor maturation along developmental trajectories, suggest that - just like genetic determinism manifests itself at the tissue and organ scales - the acquisition of multilineage competence occurs through “lineage maturation” as multipotent progenitors change through rounds of cell divisions.

## Materials and Methods

### Animals

Animal care and experiments in this study were conducted in accordance with the Animal Welfare Guidelines of the NIH and the Laboratory Animal Guidelines of China. Adult *Ciona robusca* were purchased from M-Rep, USA or collected from coastal waters in Rongcheng, China, maintained in artificial seawater with constant lighting, and used for experiments within one week of arrival.

### Sample Barcoding(SBC) Plasmids

To generate reporter transcripts with unique tags detectable by the 10X Chromium Single Cell 3’ Gene Expression Profiling System, 9 bp-short tags were inserted into *Mesp>GFP* construct 276 bp after the stop code of the EGFP coding region between the KpnI and XcmI restriction sites. The position of the tags on the 3’UTR of the transcripts is optimized to ensure effective detection by RNA sequencing.

### Sample Multiplexing

Samples from different developmental stages or CRISPR knock-out conditions were electroporated with a pair of sample barcoding (SBC) constructs as a unique label for the multiplexing strategy.

Detection of SBC in pairs (as designed) is the basis for determining the origin of each cell. The best practice to avoid technical variation between samples is to prepare the fertilized eggs for all samples in a large pool and electroporate the paired SBCs individually at one-hour intervals according to the experimental design. In this study, 10 batches of embryos of 5-14 hpf were harvested once and pooled together for cell suspension preparation. Mesp enhancer driven SBC reporters were used for barcoding and FACS enrichment of cardiopharyngeal lineage cells, while the universally expressing Eef1a enhancer-driven SBC reporters were used to multiplex whole embryo time series samples.

### Single-cell suspension & fixation

For both TVC trajectory and whole embryo data sets, single cell suspensions were prepared as described^68^. Embryos and larvae at the desired developmental stages and experimental conditions were collected in 5 mL borosilicate glass tubes (Fisher Scientific, catalog no. 14-961-26) and washed with 2 mL Calcium- and Magnesium-free artificial seawater (CMF-ASW: 449 mM NaCl, 33 mM Na2SO4, 9 mM KCl, 2.15 mM NaHCO3, 10 mM Tris-Cl at pH 8.2, 2.5 mM EGTA). Embryos and larvae were dissociated in 2 mL of 0.2% trypsin (w/v, Sigma, T-4799) CMF-ASW by pipetting with glass Pasteur pipettes. Dissociation was stopped by adding 2 mL of filtered ice-cold 0.05% BSA CMF-ASW. Dissociated cells were passed through a 40 µm cell strainer and collected in 5 mL round-bottomed polystyrene tubes (Corning Life Sciences, ref. 352235). Cell suspensions were transferred to 2.0 mL LoBind Tube (Eppendorf, Cat. No.022431102) and collected by centrifugation at 800 g for 3 minutes at 4°C, followed by two washes with ice-cold 0.05% BSA CMF-ASW. After dissociation, cell suspensions were again concentrated into a pellet at 800 g for 3 minutes at 4°C, and most of the supernatant was removed, leaving over 100 µL. The pellet was then thoroughly resuspended by gentle pipetting. Prechilled 900 µL −20°C methanol was added drop-wise to the cell suspension while gently tapping the tube to avoid aggregation. When all 900 μL of methanol was added, the tubes were thoroughly mixed by inversion 3 times and placed on ice for 30 minutes (inversions of 10 minutes each). Cell suspensions can be stored at −80°C for weeks without significant change in RNA integrity.

### Cell Rehydration and Immunostaining for FACS

Fixed cell suspensions were removed from the −80°C freezer and allowed to settle on ice for 10 minutes. Cells were harvested by centrifugation at 1500 g for 5 minutes at 4°C to avoid cell loss, followed by two washes with ice-cold 3X SSC high BSA rehydration cocktail (SSC: 3X, BSA, 0.25%, DTI, 40mM). The cell suspensions were then concentrated to a pellet at 800 g for 3 minutes at 4°C. After removing 300 µL of supernatant, the cell pellets were carefully and gently resuspended in the remaining 200 µl of 3X SSC high BSA rehydration cocktail. Alexa-488-conjugated GFP antibody (BioLegend, FM264G) was added to the cell suspensions at 5 µL to 200 µL dilution, and then incubated at 4°C with gentle agitation for 2 hours. After incubation, cells were retrieved by centrifugation at 800 relative centrifugal force (rcf) for 5 minutes at 4°C and then resuspended with 500 µL of ice-cold 3X SSC high BSA rehydration cocktail, followed by gentle agitation for 10 minutes. The cells were washed three times with 3X SSC low BSA rehydration cocktail (SSC: 3X, BSA, 0.05%, DTI, 40mM) and used for Alexa 488 positive sorting within 1 hour.

### Whole embryo barcoding

Hashtag barcoding antibody-oligos (HTOs) were conjugated to the mouse anti-dpERK monoclonal antibody (Sigma-Aldrich, M9692) using the CITE-seq hyper antibody-oligo conjugation protocol (https://cite-seq.com/), and the antibody was incubated with cell suspensions in the whole embryo study using the same immunostaining procedure as for the Alexa-488-conjugated GFP antibody. However, the information obtained from the mouse anti-dpERK monoclonal antibody is not discussed in this study.

### 10X Single Cell 3’ Gene Expression library preparation & sequencing

Rehydrated cell suspensions from whole embryos or FACS-purified B7.5 lineage were collected at 800 relative centrifugal force (rcf) for 5 minutes at 4°C. The supernatant was removed as much as possible before adding 33.4 µl RT mix (10X) + 46.6 µl water = 80 µl to the bottom of the tube. Gently pipette ∼ 30 times to ensure cells are thoroughly resuspended. The 10X Single Cell 3’ Gene Expression Library was prepared according to the 10X protocol except that 1 µl 0.2 µM of SBC additive primer (5’-ttgccgctatttctctgggtacc −3’) was added to the 10X cDNA amplification mix to generate a separate SBC library in parallel with the gene expression library. The 3’ gene expression library was purified using the standard 10X protocol, while the SBC library was purified using two rounds of 2X SPRI from the supernatant of the 0.6X SPRI 3’ gene expression library purification step.

### Single Cell Sequencing data preprocessing

20 SBC barcodes including the flanking sequences were added as pseudogenes to the *Ciona robusca* reference genome (http://ghost.zool.kyoto-u.ac.jp/download_ht.html). Raw sequencing data was mapped to the reference genome using 10X Genomics’ Cell Ranger (Version 3.0.1) pipeline to generate UMI count matrix for the downstream analysis. An SBC counts matrix was generated individually using Bowtie.

### Imputation of developmental time

We imputed developmental time of cells lacking a barcode-derived developmental age by using an approach inspired by ancestry voting [1,2]. The dataset was split into cells with and without time stamp based on electroporated barcodes. The twenty nearest neighbors for each cell without a time stamp were identified. The inferred developmental time of a time stamp was set to the median of the time stamps of its k-nearest neighbors. Only in cases where the standard deviation of the developmental time of the k-nearest neighbors was below 1 did we assign an inferred time stamp to a cell. Using all cells with a timestamp as a ground truth and performing this label transfer approach for each cell with time stamp individually, we estimate that we infer the correct transcriptional age in 80.4% of the cases and are within 1h of development in 95.9% of cells. Thus, only 4.1% of the cells might have gotten a severely wrong inferred timepoint, in addition to the 20.3% of cells that did not get any time label assigned. For the final developmental time, we first considered if a cell had a barcode-derived time stamp, in case of differences between the inferred and barcode-derived developmental time, the barcode-derived developmental time was retained.

### Mesenchymal trajectories

To obtain developmental trajectories of individual endodermal and mesenchymal lineages, we subclustered the whole mesenchyme and endoderm independently and grouped the resulting higher resolution Leiden clusters by expression of marker genes and continuity of the UMAP projection. We constructed a UMAP for each isolated branch and calculated a branch specific pseudotime using Scanpy, setting a random cell with the lowest barcode-derived developmental time stamp as root cell.

### Mesenchymal trajectories

To obtain developmental trajectories of individual endodermal and mesenchymal lineages, we subclustered the whole mesenchyme and endoderm independently and grouped the resulting higher resolution Leiden clusters by expression of marker genes and continuity of the UMAP projection. We constructed a UMAP for each isolated branch and calculated a branch specific pseudotime using Scanpy, setting a random cell with the lowest barcode-derived developmental time stamp as root cell.

### Gene clustering

Gene clusters were calculated by transposing the gene count matrix, filtering out genes that were expressed in less than 3 cells in the derived matrix and performing dimensional reduction and leiden clustering on the transposed matrix, similar to the standard procedure in the identification of cell clusters. Mature gene clusters were defined as clusters of genes that were upregulated shortly before and around the time of lineage bifurcation, while immature gene clusters were defined as the first gene cluster that was upregulated in the founder cell population.

### Marker gene identification

A Wilcoxon rank sum test was used to identify marker genes for different cell states. To enrich for significant marker genes, we filtered the putative markers for an average log2FC of 0.25 and an adjusted p-value < 0.1.

### sgRNA design and library construction

#### CRISPR library construction

All sgRNAs were cloned into the *2mU6>sgRNA(F+E)* scaffold as per Stolfi, et al.^49^ *2mU6>sgRNA(F+E)* plasmids were digested with BsaI-HF (NEB R3535, discontinued) in rCutsmart buffer for optimal positivity rates. Oligos ordered from Sigma at 50 mM concentration in water, were annealed by combining 10 uL of forward and reverse oligos and incubating in a thermocycler with the following protocol: 1) 95 °C for 30 seconds (melt) 2) 72 °C for 2 minutes, 3) 37 °C for 2 minutes, 4) 25 °C for 2 minutes. Annealed oligos were diluted 1:50 ul in TE buffer, and ligated into an opened vector using Instant Sticky-end Ligase Mix (NEB M0370S) with 0.5 µL vector, 2.5 µL diluted oligo, 3 µL ligase mix. 3 µL of ligation mix were transformed into 15 µL Stellar Competent cells (Takara 636763) and plated on AMP LB selection plates followed by overnight incubation at 37 °C. 4 colonies per plate were selected for colony PCR using the U6-F-910 forward primer (5’-caattgccccaagctctcttc-3’) and the guide RNA specific sgRNA-R oligo (same as the R for annealing diluted to 10 µM). Positive colonies were grown overnight in 4 ml LB amp overnight, miniprepped, and sent for sequencing using the seqU6-F primer (5’-ggatcgcgcgagccc-3’). Sequence results were checked to assure no mutations were present in the guide RNA sequence and correct clones were frozen down in Hogness Freezing Medium (Bioworld 30629174-1) and stored at −70 °C.

### CRISPR screen data collection and analysis

For each gene in the screen, the following electroporation conditions were used: Mesp>NLS::LacZ (nuclear marker) - 20 µg, Mesp>hCD4::mCherry (cell membrane marker) - 20 µg, Mesp>NLS::Cas9::NLS - 25 µg, 2mU6>sgRNA.1.2.3 - 80 µg. The total DNA amount used for electroporation did not exceed 150 µg. Embryos were incubated at 18 °C for 15 hours post fertilization (hpf). At 15 hpf embryos (approx. FABA stage 25) were fixed in 4% paraformaldehyde and were used for fluorescent in situ hybridization as described ^53^ using an intronic Tbx1/10 digoxigenin-labeled RNA probe^36^. 3 biological replicates were performed for each condition. 3D stacks were collected on the Leica WLL SP8 Confocal microscope using a 63x oil immersion lens. Images were analyzed using Imaris Bitplane software for presence of *Tbx1/10* nascent transcripts, cell division orientation along the anterior-posterior and medio-lateral axis, and for asymmetric cell division as determined by cell size post division. In total over 1,500 embryo halves were analyzed. Graphs and statistical analysis performed in Python using Matplotlib and Seaborn pipelines.

### Depdc Enhancers Characterization

*Depdc1b* enhancers were predicted based on the TVC-specific chromatin accessibility identified by Racioppi et al^54^. Three versions of the enhancers (3691/-1, −2537/-1 and −1969/-1) including the native promoter of the *Depdc1b-KY.Chr2.2230* gene were amplified using the following primers

(5′-tccgcgcgccGTATIGGGGAACCCGGGATAAAAATATIGAAACGTC-3′ and

5′-accgcgccgcTCTIATGTGTITGATAAACCTGTIAAACGAAAAGC-3′ for −3691/-1,

5′-ttagcgcgccAACAATATIGTCCTGGTITGACTITC-3′ and

5′-accgcgccgcTCTIATGTGTITGATAAACCTGTIAAACGAAAAGC-3′ for −2537/-1,

5′-gcgcgcgccTGCTACTIATAGGAAGAAATATATAGTITITG-3′ and

5′-accgcgccgcTCTIATGTGTITGATAAACCTGTIAAACGAAAAGC-3′ for −1969/-1) with the addition of AscI and NotI restriction sites. The enhancers were subcloned into the AscI-NotI restriction site of the 2xGFP vector. The enhancer (−2786/-2062) was amplified using the following primers (5′-tccgcgcgccCATIATIACGACTCTAAAAGAGCGCTGATAATIGTIAAACGC-3′ and

5′-gatctagaTCTGCTATATACGTCGGCCCTGTGGCAGGTGGATICTIAACAT-3′) cloned into the AscI-XbaI restriction site of bpfog: :2xGFP vector. Using the −3691/-1>2xGFP reporter as a template, ΔFlank, ΔGata4/5/6, Δ(Flank+Gata4/5/6), and ΔFoxa1/2/3 were amplified using the Mut Express II Fast Mutagenesis Kit V2 (Vazyme, Cat. No. C214-01) with the following primers

(5′-TIGCATGGCCGTGCGAGATAAGGAGCCATIACGG-3′ and

5′-CTCGCACGGCCATGCAAGTCTATAGGATACAGAGGCC-3′ for ΔFlank,

5′-TIGGAGTIAATIAGCCATIACGGTCGTCTGTGTITACTG −3′ and

5′-GTAATGGCTAATIAACTCCAAGTGACGGCCATGCAA −3′ for ΔGata4/5/6, 5′-

TIGCATGGCCGTAGCCATIACGGTCGTCTGTGTITACTG-3′ and 5′-

GTAATGGCTACGGCCATGCAAGTCTATAGGATACAGAGGCC-3′ for Δ(Flank+Gata4/5/6), 5′-

TIACGGTCGTCTCTGATGTITAGAAACAGGTIGACTGGGGG-3′ and 5′-

AAACATCAGAGACGACCGTAATGGCTCCTIATCTCGCAATIAACTC-3′ for ΔFoxa1/2/3) and cloned into the AscI–NotI restriction site of the 2xGFP vector.

### Motif enrichment analysis

#### Motif Selection

Motifs were selected from Cis-BP^58^, Aniseed^69^, and HOMER^70^. Motif database is available here: https://github.com/kewiechecki/CrobustaTFs. For each unique transcription factor, the PWM with lowest Shannon entropy was selected. Motifs were searched in the C. robusta accessome^54^ using [motifmatchr](https://bioconductor.org/packages/release/bioc/html/motifmatchr.html) with default parameters.

#### Motif Accessibility

ATAC-seq reads were mapped to each motif site using bedtools. Differential accessibility was calculated using DESeq2. Motifs were considered differentially accessible if the likelihood ratio test returned a false discovery rate > 0.1.

DA motif sets were defined as follows:

Motif Set log2 fold change cutoff
Open in FoxF: FoxF_KO vs. control > 1
Closed in FoxF: FoxF KO vs. control < −1
TVC accessible mesp_dnFGFR vs. control < −1 or mesp_MekMut vs. control > 1
ATM accessible mesp_dnFGFR vs. control > 1 or mesp_MekMut vs. control < −1
ASM accessible handr_dnFGFR vs. control < −1 or handr_MekMut vs. control > 1
Heart accessible handr_dnFGFR vs. control > 1 or handr_MekMut vs. control < −1

#### Motif Annotation

Motif sites were assigned to a gene if they overlapped the gene body, promoter, or distal region of a gene. The 5’ UTR, coding regions, introns, and 3’ UTR were considered the gene body. −1107 to +107 bp from the TSS was considered the promoter. −10 kbp from the TSS or +10 kbp from the TIS were considered distal elements.

#### Enrichment

All enrichments were calculated using a hypergeometric test for occurrence of a motif in the target motif set vs. occurrence in the background. I tested for motif enrichment in both sets of differentially accessible peaks in each condition, all peaks annotated to a gene cluster, and TVC accessible peaks annotated to a gene cluster. I also tested for enrichment of peaks annotated to specific feature types. We ran separate tests using all accessible motifs as a background and using TVC-accessible peaks as a background.

## Authors contributions

Y.B. performed the CRISPR/Cas9 screen on candidate mature genes (Figure 4) and coordinated with L.C. and other authors to assemble the final manuscript.

A.K. performed the work on cell cycle progression coupling with *Tbx1/10* and *Depdc1b* expression (Figures 1, 6 and 7), and helped coordinate the assembly of the manuscript.

A.T. performed the initial computational analyses of scRNA-seq data (Figures 2 and 3), including the establishment of ShinyApps.

J.B. performed the complementary analyses of the whole embryo scRNA-seq dataset (Figure 3).

P.Z. performed the cis-regulatory analysis of *Depdc1b* enhancers (Figure 6).

K.W. performed the motif enrichment analysis (Figure 5), and helped quantify the *Depdc1b^CRISPR^* phenotypes (Figure 4).

N.K. performed the analysis of the impact of *Depdc1b^CRISPR^* on oriented and asymmetric division and *Tbx1/10* expression.

M.F. quantified the Depdc1b^CRISPR^ phenotype using morphometric measurements (Figure 4).

M.B. and O.M. assisted Y.B. with the analysis of CRISPR/Cas9-mediated mutagenesis of candidate “mature” genes (Figure 4).

W.W. performed all scRNA-seq experiments, including the design of barcoding reporters and optimization of rehydration (Figures 2, 3), and oversaw the work on the *Depdc1b* enhancer (Figure 6)

L.C. designed the study, acquired funding, oversaw work and progress, coordinated data collection and manuscript preparation, and wrote the paper.

## Acknowledgements

We are grateful to Drs. Alberto Stolfi and Basile Gravez for preliminary work on *Depdc1^CRISPR^*, and cell cycle progression and coupling with gene expression, respectively. This work was supported by NIH awards R01 HL108643 and R01 HD096770 to L.C, and by core funding to the S16 (Christiaen) group at the Michael Sars Centre, from the University of Bergen and the Research Council of Norway.

---

**Figure 1-S1.**
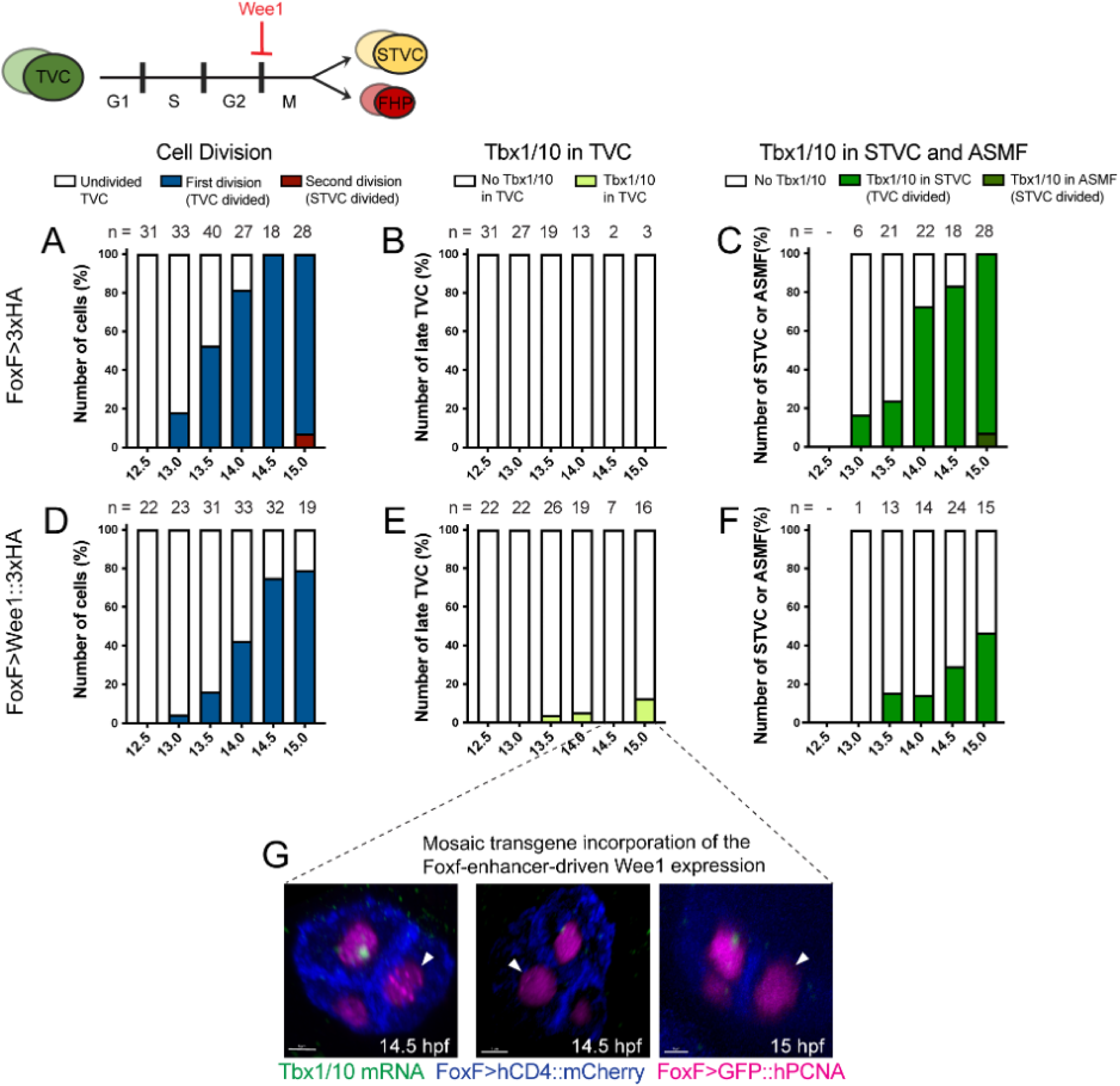
Inhibition of mitotic entry in TVC lineage resulted in reduced Tbx1/10 expression in STVC. **A-F.** Cell division patterns (A and D) and Tbx1/10 expression upon cell mitosis suppression perturbations (Wee1::3xHA) in the late TVCs (B and E) and STVCs (C and F). Note that the results of A, B, D, E used the same dataset as Figure 1F, 1H, 1F’, and 1H’. **G**. Confocal images showing mosaic transgene inheritance of Wee1::3xHA under the Foxf enhancer in some samples. Magenta: nuclei (GFP::PCNA); Blue: cell membranes (hCD4::mCherry); White arrowhead: late TVC, Scale bar = 5mm.

**Figure 1-S2.**
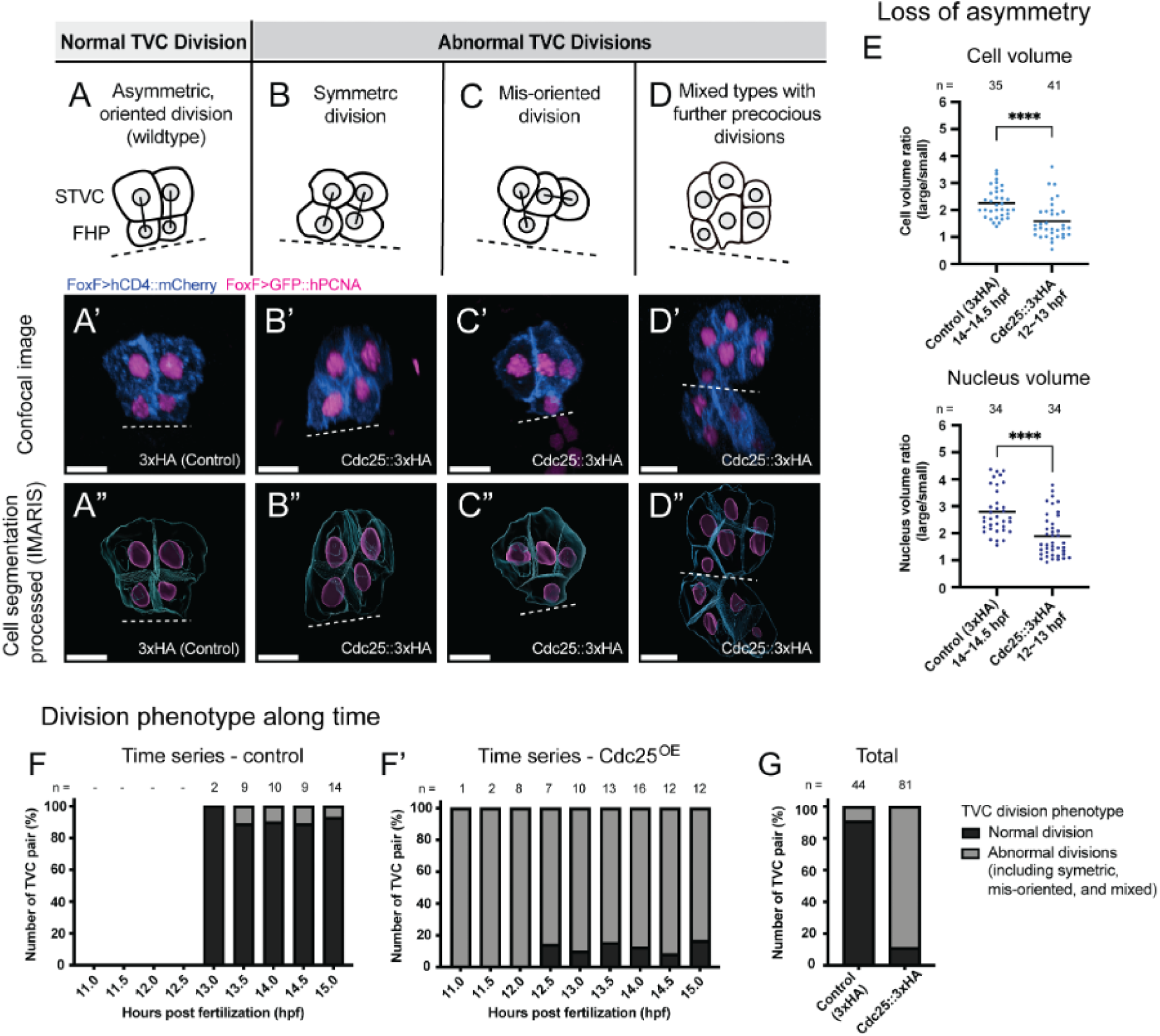
Cdc25-overexpressing TVCs undergo abnormal cell division with the loss of asymmetry and loss of medio-lateral orientation. **A-D.** Schematic diagrams of four categories of normal and abnormal division phenotypes of TVC. Normal division (A, asymmetric oriented) produces a pair of lateral large STVCs and small medial FHPs. Abnormal divisions include symmetric (B), mis-oriented (C), and the mixed types (D). **A’-D’** and **A”-D”.** Representative images of each division category under the control and mitosis perturbations (Cdc25) conditions. Magenta: nuclei (GFP::PCNA); Blue: cell membranes (hCD4::mCherry); Dashed line: embryo midline, Scale bar = 10mm. **E-F**. Division phenotype of TVC throughout developmental time points in the control (E) and Cdc25-overexpressing (E’) samples. With results from different time points combined, F shows the total effect of Cdc25 on mis-regulating mitosis. **G.** Quantification of the cell volume ratio (left) and the nuclear volume ratio (right) of TVC descendant cells in the control and Cdc25-overexpression conditions. **** P < 0.0001.

**Figure 1-S3.**
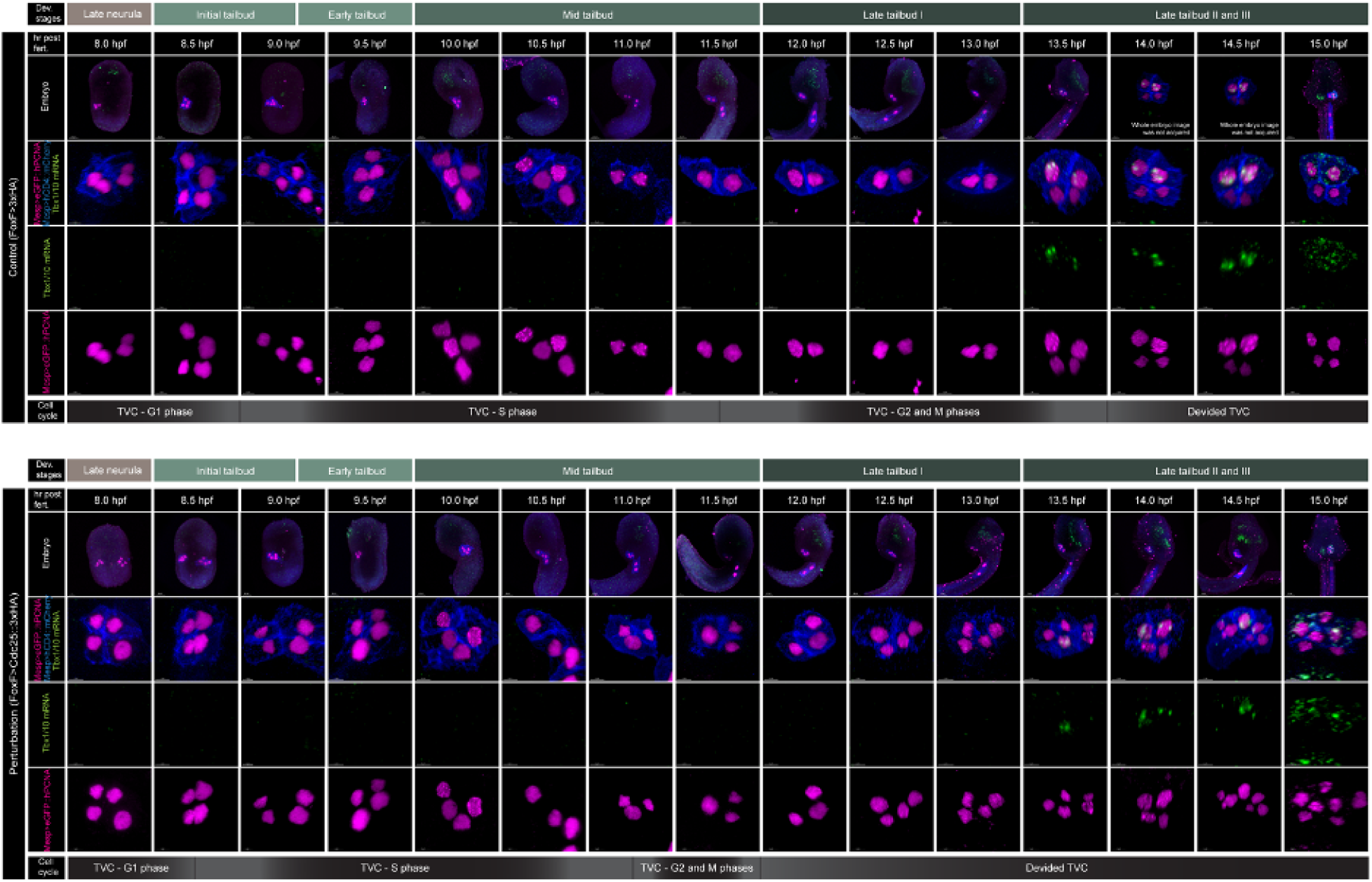
Representative confocal images of Cdc25^OE^ perturbation time series. The division and Tbx1/10 expression phenotypes can be seen in the control (Top) and the Cdc25-overexpressing (Bottom) samples. Embryogenesis developmental stages and the cell cycle details of TVC are shown. Magenta: nuclei (GFP::PCNA); Blue: cell membranes (hCD4::mCherry); Scale bar = 20mm for embryos, 5mm for enlarged images.

**Figure 1-S4.**
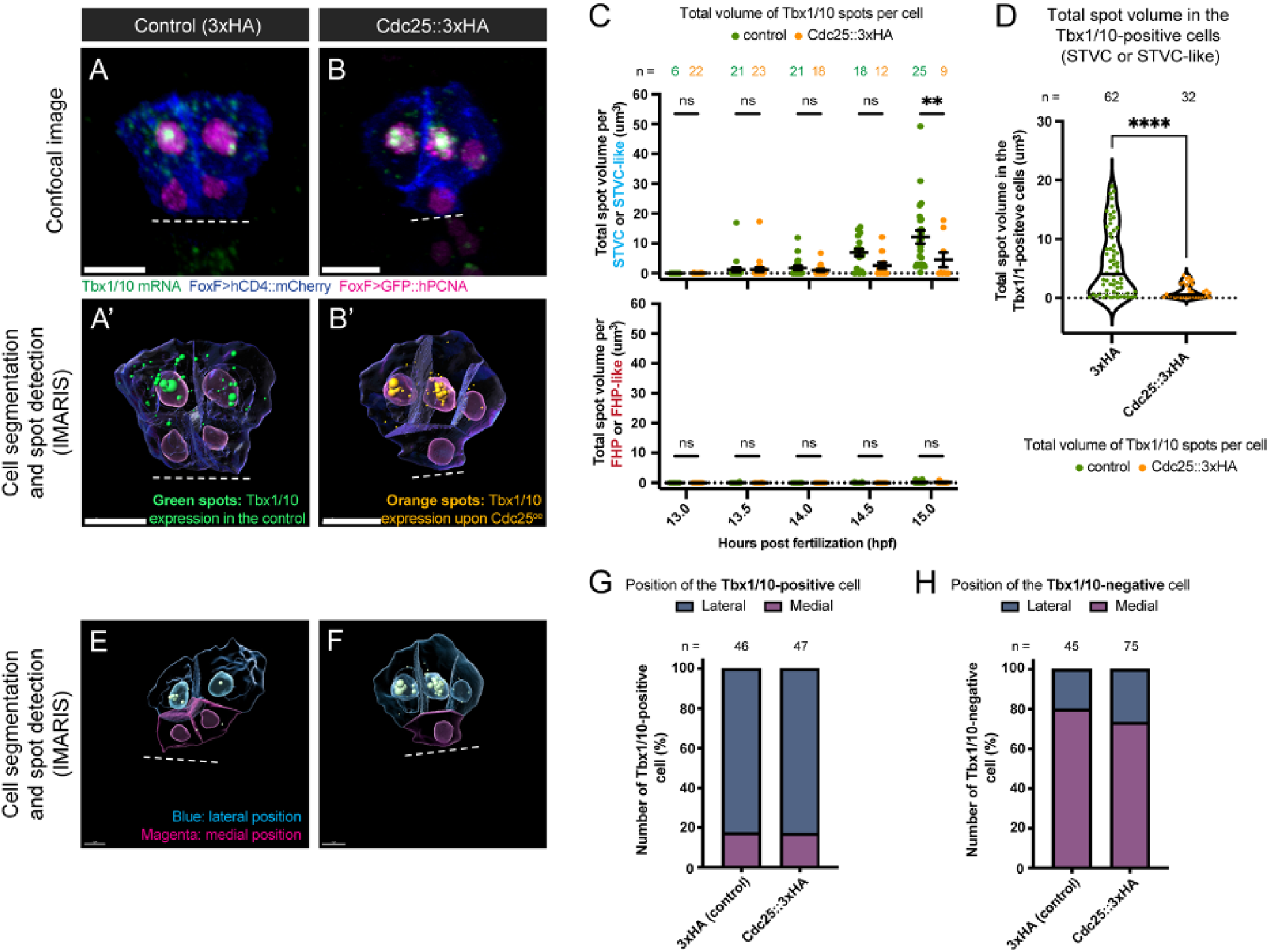
The accumulation of Tbx1/10 in the TVC descendants is affected by the mitotic perturbation of Cdc25, with niche-driven mechanisms involved. **A-B** Representative confocal images of Tbx1/10 expressing in TVC descendants in the control and Cdc25^OE^ samples. Magenta: nuclei (GFP::PCNA); Blue: cell membranes (hCD4::mCherry); Dashed line = embryo midline; Scale bar = 10mm. Note that panel B uses the same image data as Figure 2S panel C for representation. **A’-B’**. Processed images with cell segmented and Tbx1/10 FISH spot signal detected in STVC or STVC-like cells. Green spot: FISH signal detected in the control; Orange spot: FISH signal detected in the Cdc25^OE^ cells. **C.** Developmental quantification of total volume of spots detected in individual STVC(-like) cells (top) and individual FHP(-like) cells (bottom) under the control and Cdc25^OE^ conditions. ns, not significant; ** P < 0.01. **D.** Total spot volume in Tbx1/10-expressing cells under the control and Cdc25^OE^ conditions. **** P < 0.0001. **E-F** and **E’-F’**. Representative confocal images of TVC descendants in the control and Cdc25^OE^ samples. Images are processed by cell segmentation, FISH spot detection, and labeled by colors based on cell localization. Processed blue cell: lateral-most localization; Processed magenta cell: median localization. Scale bar = 5mm. **G-H**. Localization analysis of Tbx1/10-expressing (G) and Tbx1/10-negative (H) cells under the control conditions and Cdc25 perturbations.

**Figure 1-S5.**
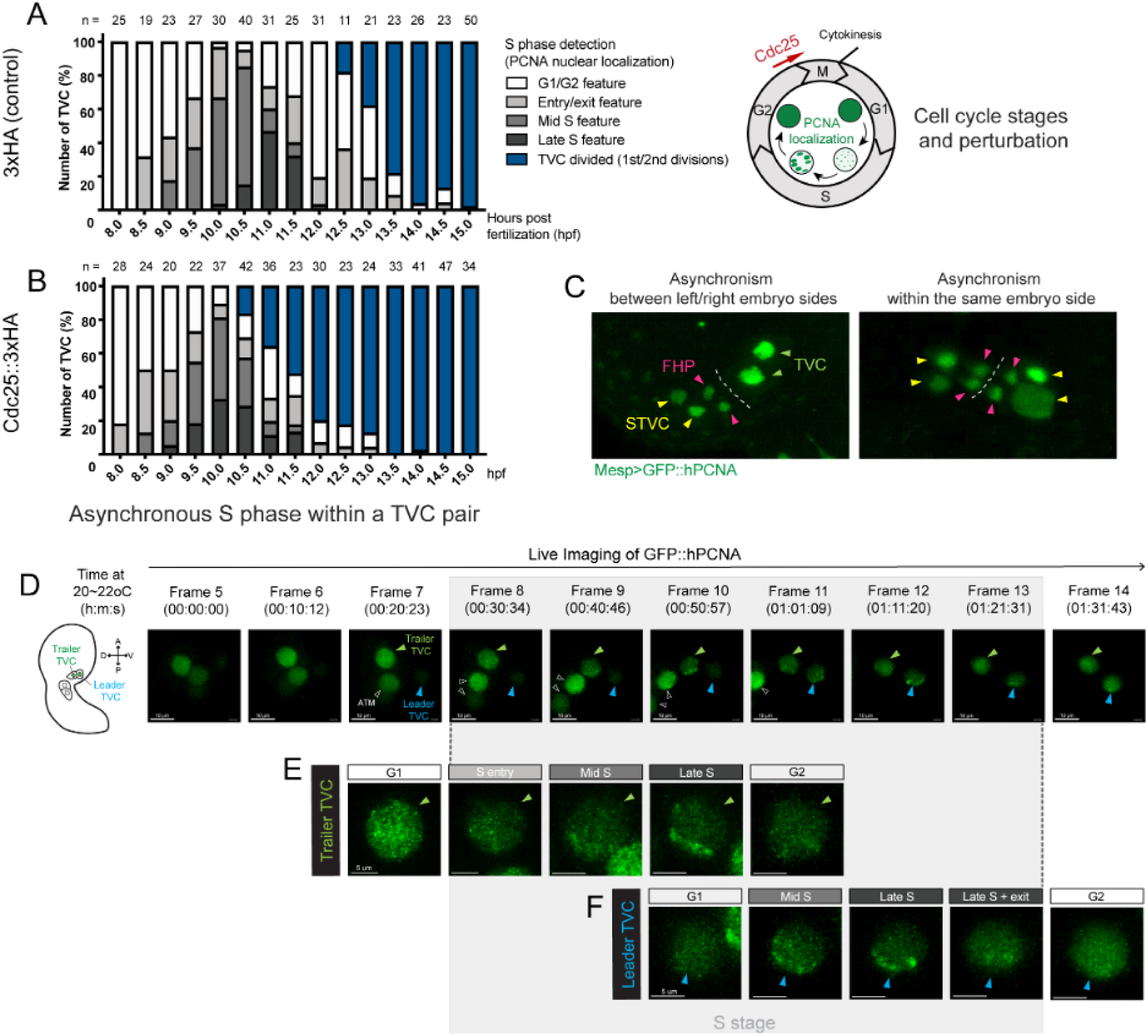
Cell cycle progression dynamics of TVCs under perturbations and in nature. **A.** Schematic of cell cycle stages, genetic perturbation (Foxf>Cdc25) of G2/M transition, and variability of PCNA puncta size in the TVC nuclei associated with cell cycle progression. **B-C**. Developmental distribution of S phase (four PCNA localization patterns) and cell division of TVC in the control and Cdc25^OE^ samples. Cdc25-overexpressing TVCs are permitted to divide right after passing through S/G2 transition (C). **D**. Confocal images showing asynchronous division of TVC-lineage cells. (Left) Asynchronous TVC division between left/right body sides. (Right) Asynchronous division within the same body side. Green: nuclei (GFP::PCNA); Green arrowhead: TVC; Yellow arrowhead: STVC; Magenta arrowhead: FHP. **E**. Live imaging of PCNA puncta dynamics in the leader and trailer TVCs along TVC cell cycle progression revealed asynchronous S phase within collective migrating TVC pair. Green: nuclei (GFP::PCNA); Blue arrowhead: leader TVC; Green arrowhead: trailer TVC; Scale bar = 10mm.

**Figure 2-S1.**
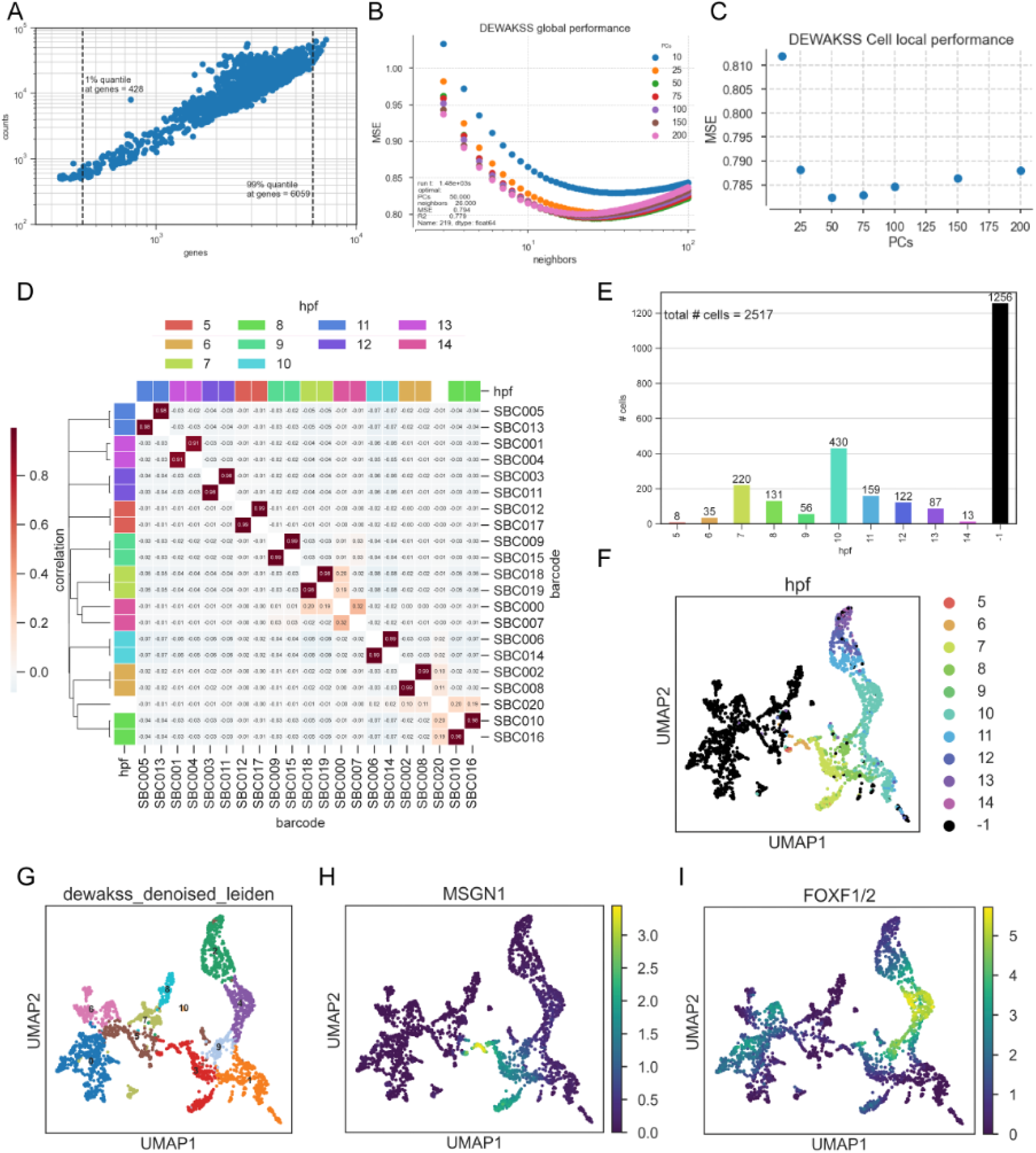
**A.** Number of genes detected vs. the number of unique molecular identifiers (UMIs) per cell. **B.** Denoising with DEWAKSS hyperparameter grid search. Each color represents the number of PCs used. X-axis is the number of neighbors, performance of DEWAKSS in mean square error (MSE) on y-axis. **C.** Performance when optimal number of PCs and neighbors per cell are used. **D.** Leiden clustering given derived kNN graph from DEWAKSS. **E.** Real-time barcodes displayed on cells clustered in D. **F-G**. TVC-lineage marker genes for expected trajectory root (MSGN1, expressed in founder cells) and branch (Foxf, expressed in cardiopharyngeal lineage. **H.** Correlations of double barcode system to assign cells to real developmental time. Pearson correlation coefficient is shown. **I.** Number of cells detected per real-time barcode pair. −1 are cells that did not pass barcode annotation threshold selection (see Materials and Methods).

**Figure 2-S2.**
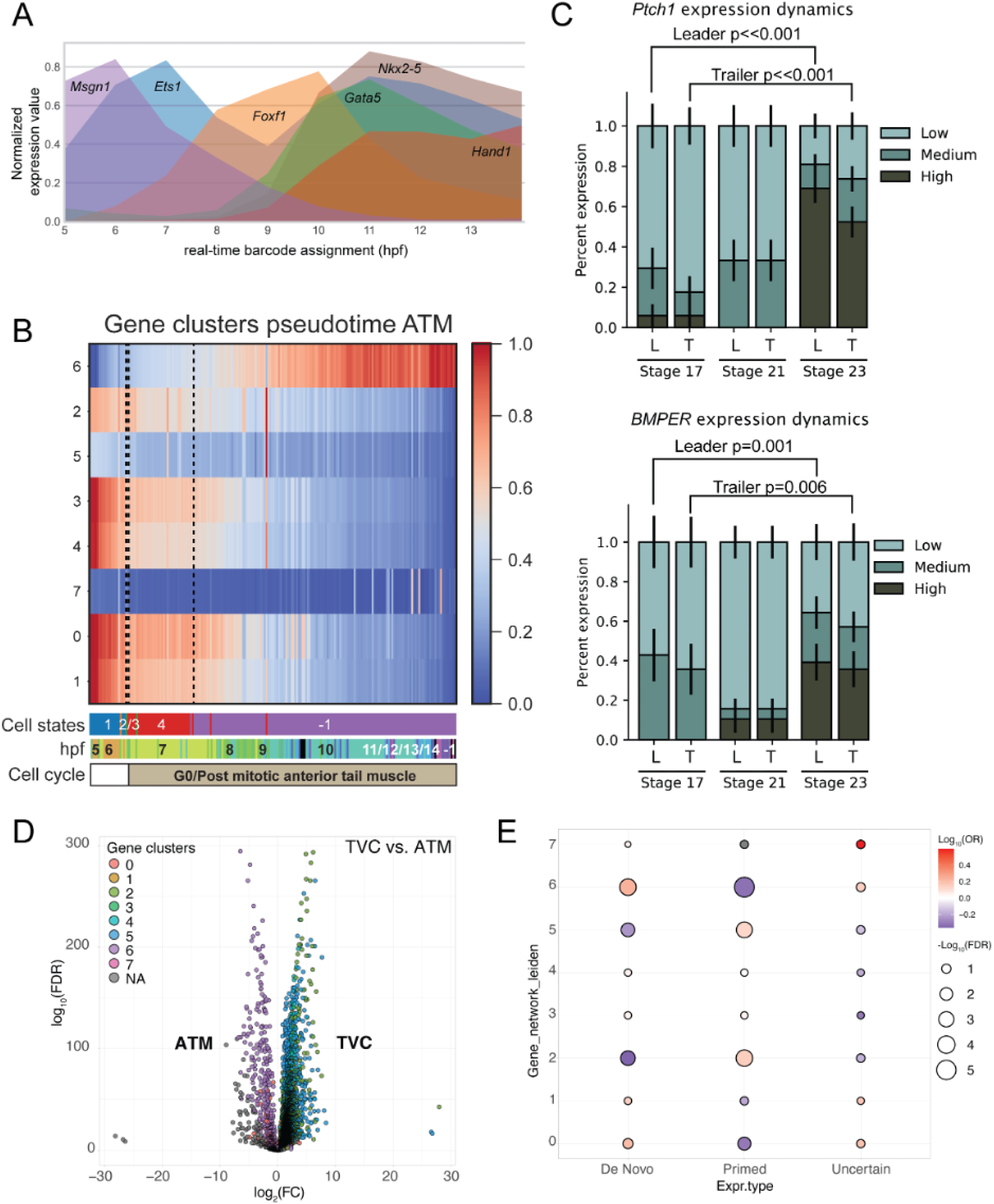
**A.** Normalized expression values of core cardiopharyngeal precursor genes (*Mesp1, Ets1,Foxf1, Gata5, Nkx2-5, Hand1*) as a function of real-time barcodes. **B.** Cell states and gene expression dynamics of post-mitotic Anterior Tail Muscles (ATMs). **C.** Dynamics of Ptch1 and BMPER expression based on FISH analysis (Figure 2L). Embryos from indicated stages are scored based on qualitative expression level (low, medium, high). Standard error of proportion is shown. P-values is determined using the Pearson’s c^2^ test. **D.** Log-fold change in TVC vs ATM gene expression grouped by identified gene clusters. **E.** Odds ratio of cluster-specific gene expression grouped by expression defined as de novo expression in STVCs, primed in the TVCs prior to division, or uncertain onset of expression with respect to TVC division

**Figure 3-S1.**
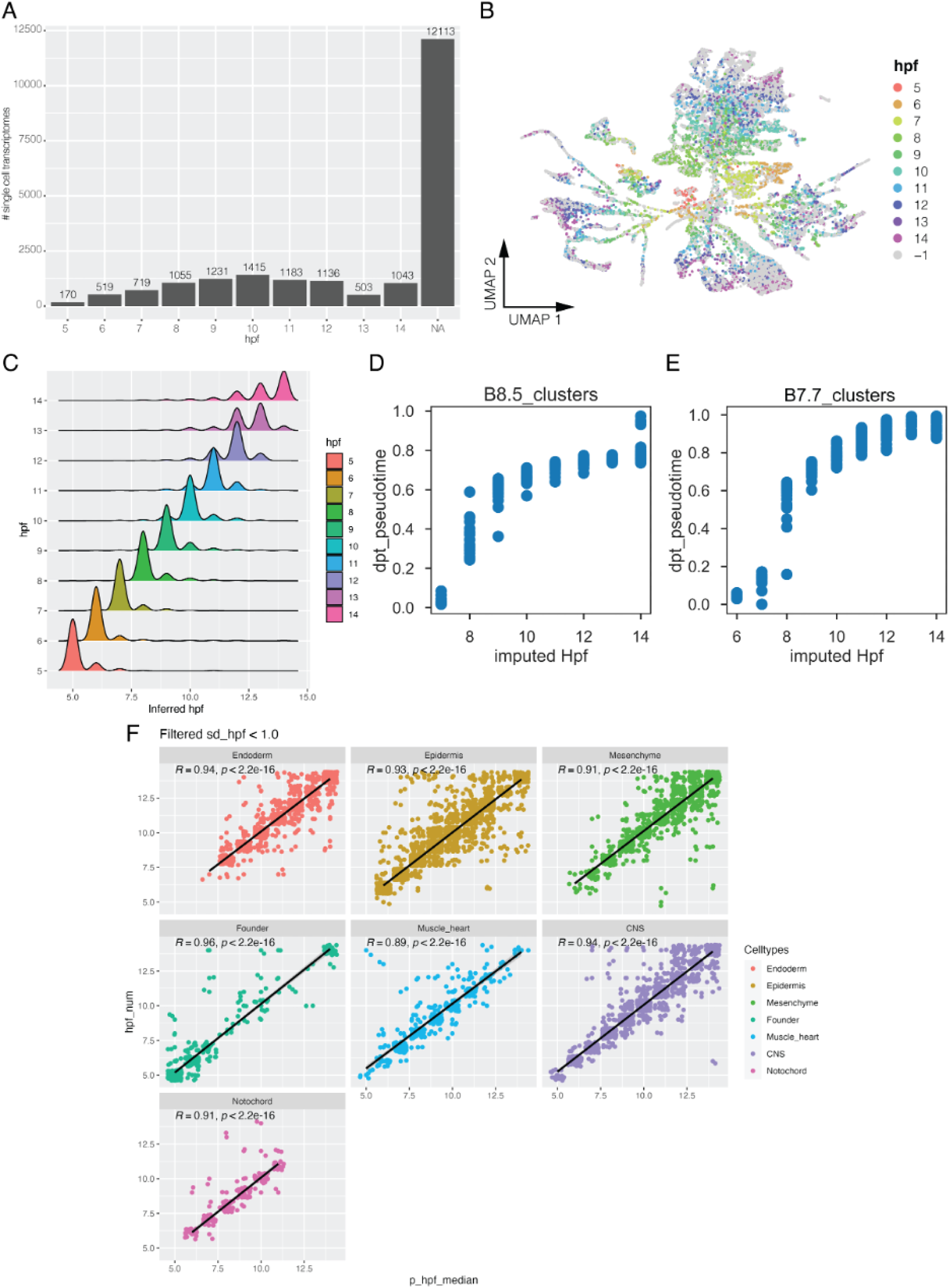
Propagation time stamps to the whole embryo dataset through label transfer. **A**. Frequency distribution of time specific barcodes across the developmental times profiled for the whole embryo dataset. Most cells did not express a time stamp. **B**. UMAP representation of the whole embryo scRNA-seq dataset labeled with barcodes-derived time stamps only, also shown in Figure3-B. **C**. Ridge plot showing the distribution of predicted developmental time through label transfer across all cells with a valid barcode-derived time stamp and binned by barcode-derived time stamp on the y axis. **D, E.** Dot plot showing the correlation of inferred developmental time with diffusion pseudotime (dpt) for the cells of the mesenchymal B8.5 lineage (D) and the mesenchymal B7.7 lineage (E). **F**. Correlation of barcode-derived developmental time with developmental time inferred through label transfer, shown for cells of each main tissue independently, as indicated in the panel.

**Figure 3-S2.**
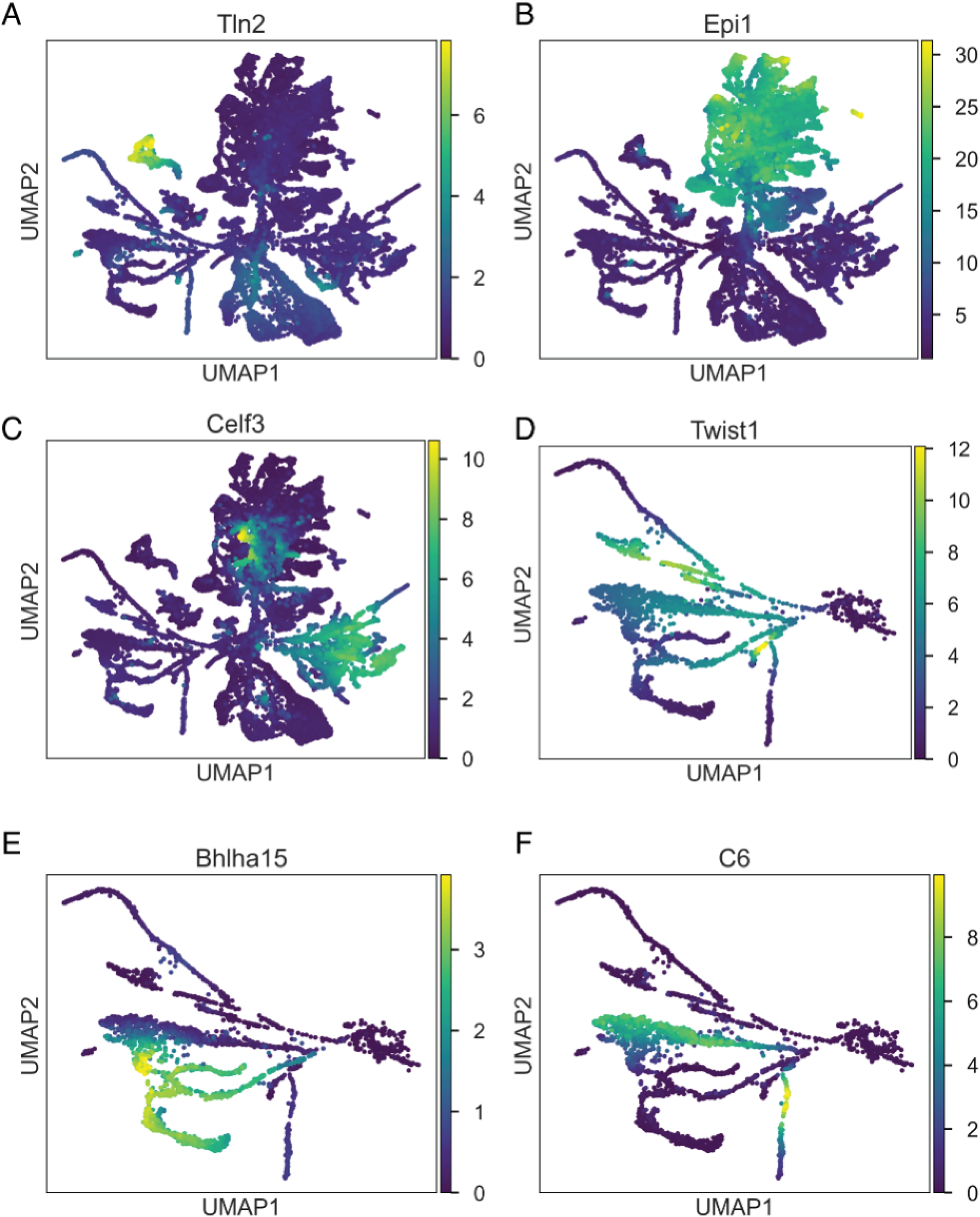
Expression of tissue-specific markers acoss the whole embryo scRNA-seq dataset. **A-C**. Expression of a notochord specific marker *Tallin* (A, KY.Chr3.779), an epidermis specific marker *Epi1* (B, KY.Chr1.2380”), a CNS specific marker *Celf3* (C, KY.Chr6.59) are shown on the UMAP representation of the whole embryo dataset. **D-F**. Expression of the A7.6 specific marker *Twist1* (D,, KY.Chr5.357), the B7.7 specific marker *Bhlha15* (E, KY.Chr3.1309), and the B8.5 specific marker *C6* (F, KY.Chr14.191), are shown on the UMAP representation of mesenchymal subset.

**Figure 3-S3.**
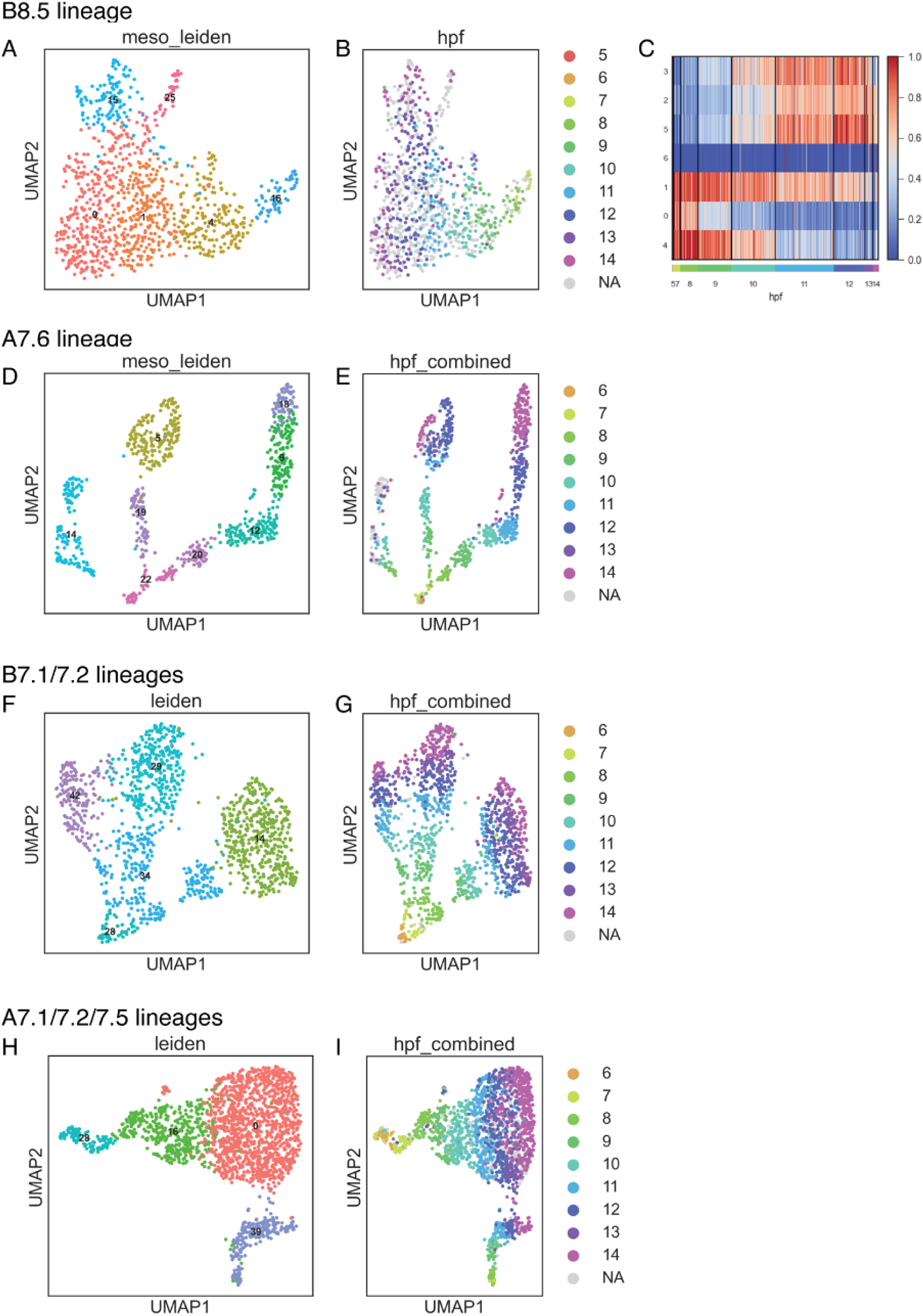
Endomesodermal trajectories. **A,B.** Differentiation trajectory of the B8.5 mesenchymal lineage colored by inferred developmental time (A) and denoised Leiden clustering (B). Note that cluster 16 corresponds to an immature state and cluster 1 to a mature transcriptomic state. **C.** Differential expression of genes during differentiation of the B8.5 mesenchymal lineage. Gene clusters 4 and 1 correspond to the immature and mature states, respectively. **D,E.** Differentiation trajectory of the A7.6 mesenchymal lineage colored by inferred developmental time (D) and denoised Leiden clustering (E). Due to the rapid specification of these lineages early in development, no immature and mature cell state could be identified at the resolution of the present dataset for the A7.6 lineage. **F,G.** Differentiation trajectory of the B7.1 and B7.2 endodermal lineages colored by inferred developmental time (F) and denoised Leiden clustering (G). **H,I.** Differentiation trajectory of the A7.1, A7.2 and A7.5 endodermal lineage colored by inferred developmental time (H) and denoised Leiden clustering (I). Note that due to the unclear clonal relationship of the trajectories in the endoderm, we did not attempt to identify compatible mature and immature states

**Figure 4-S1.**
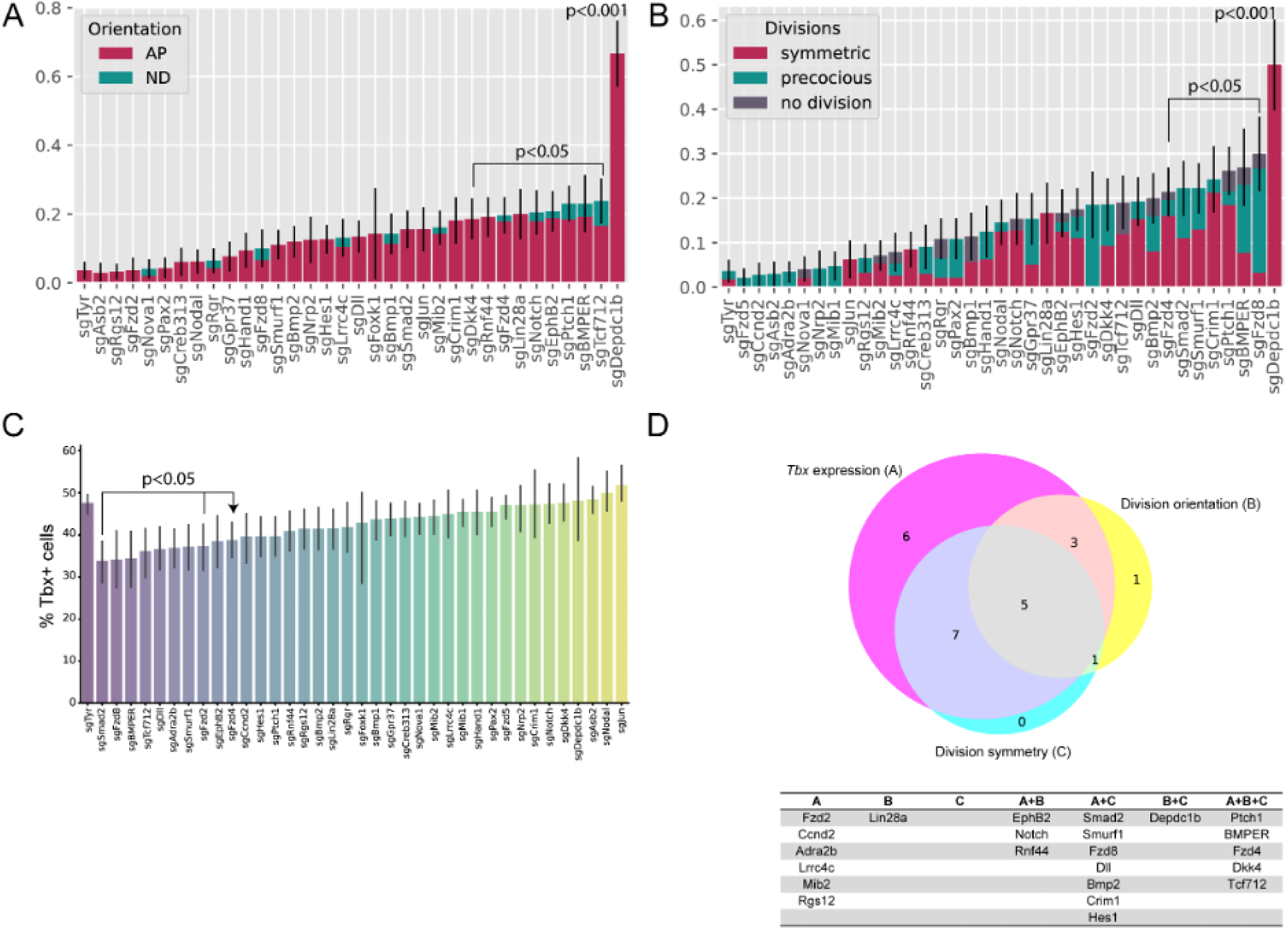
CRISPR screen for orients asymmetric TVC division. **A.** Abnormal orientation of TVC division under shown CRISPR targets. AP = Anterior-Posterior orientation, ND = No Division. **B**. Abnormal division symmetry or timing under indicated CRISPR perturbation. For both A and B error bars show standard error of proportion (SEP) for total abnormal phenotypes observed. P-values calculated using the Fisher exact test with a Bonferroni correction. **C**. Percent Tbx1/10 positive cells under indicated CRISPR perturbation. Error bars indicate 95% confidence interval. Statistical analysis performed using a Two-tailed t-test followed by Bonferroni correction. **D.** Venn diagram of top CRISPR screen phenotype hits with a table listing individual and combined phenotypes for each CRISPR condition.

**Figure 4-S2.**
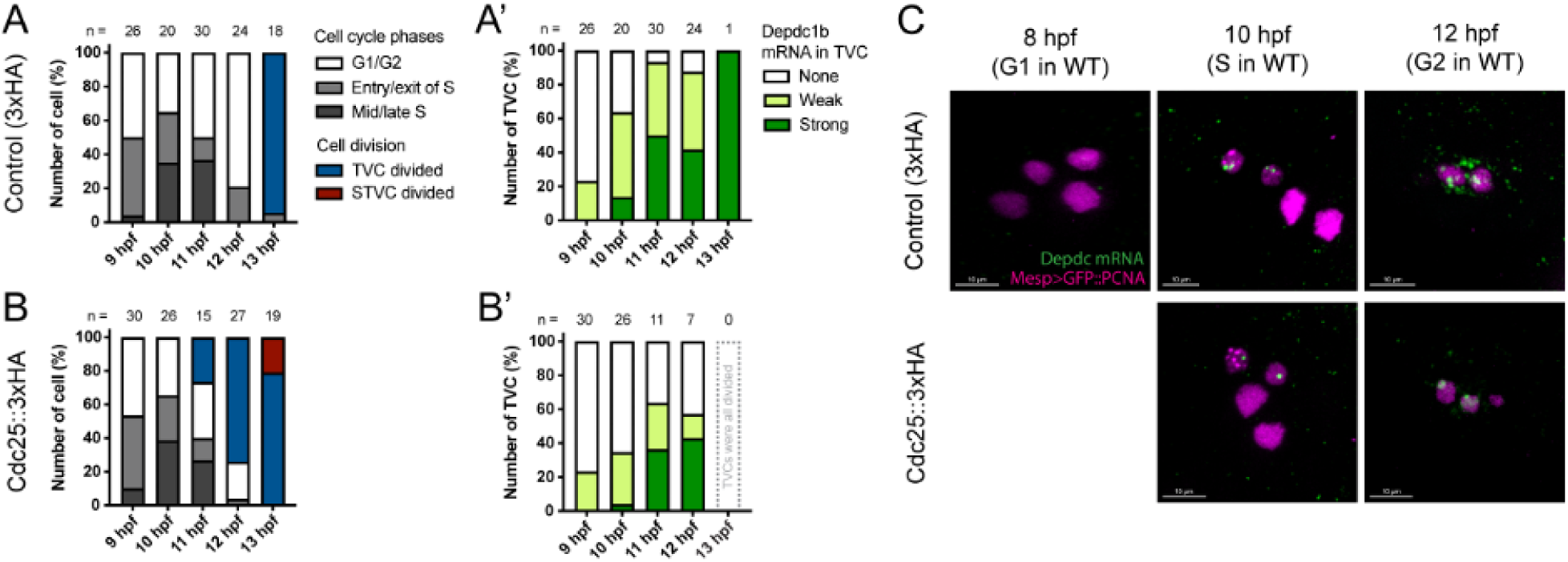
Intact G2 progression permits high accumulation of Depdc1b in TVC. **A-B.** Developmental distribution of S phase and cell division of TVC under the control and Cdc25^OE^ conditions. **A’-B’**. The corresponding Depdc1b expression phenotype along these developmental time points. Perforated bar in B’ indicates no cells can be analyzed. **C.** Representative images of Depdc1b expression at 8, 10, and 12 hpf in the control and the Cdc25-overexpressing cells. Magenta: nuclei (GFP::PCNA); Green: Depdc1b mRNA. Scale bar = 10mm.

**Figure 5-S1.**
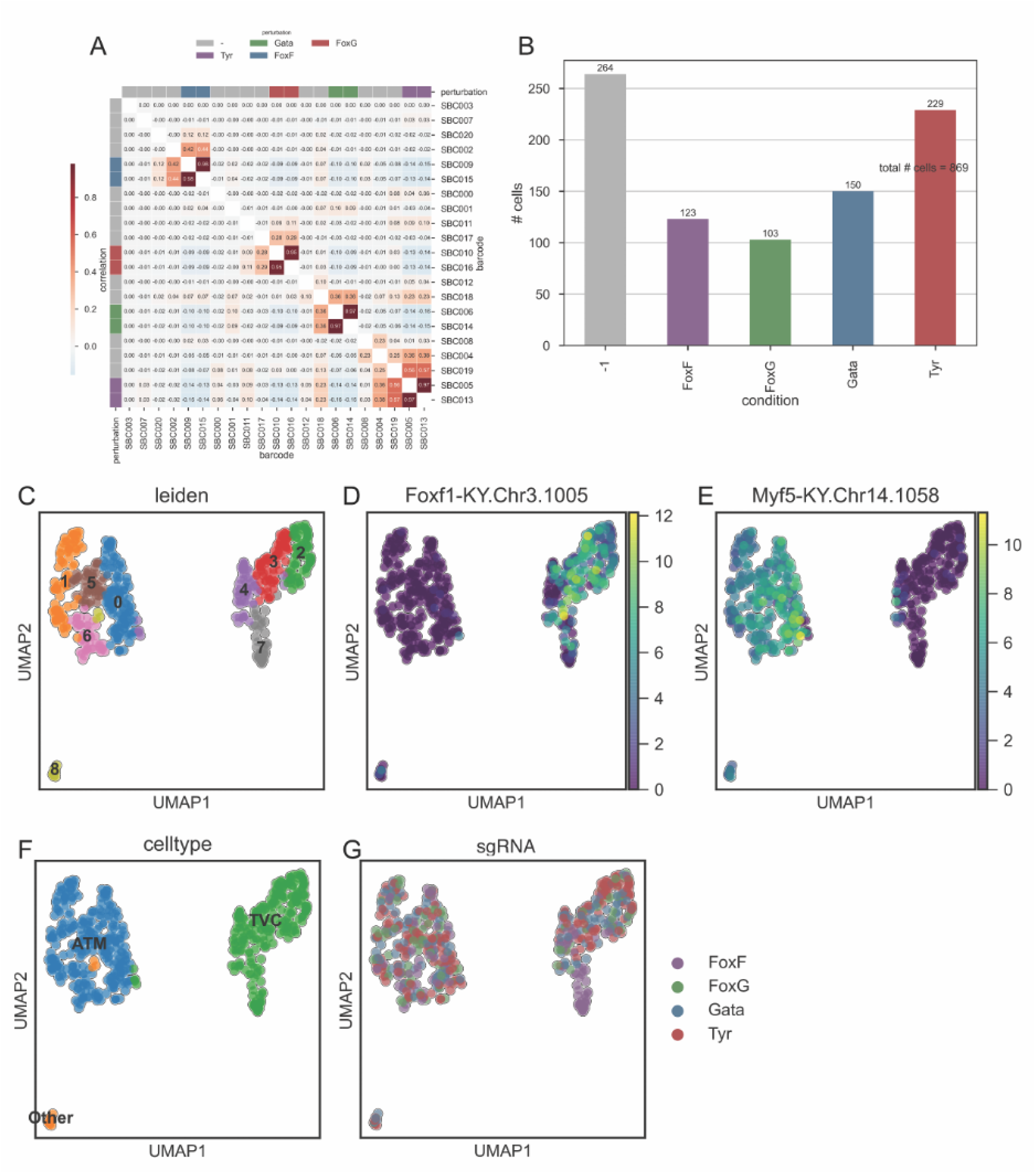
Sample barcode recovery in the CRISPR x scRNA-seq experiment. **A.** Correlation plot between co-electroporated pairs of sample barcoding constructs. **B.** Distribution of sample assignment following barcode recovery. **C**. UMAPS showing the distribution of individual single cell transcriptomes across Leiden clusters. **D-F**. Expression of key cardiopharyngeal (Foxf, D) and anterior tail muscle (Myod/Myf5, E) markers allowing annotation of Leiden clusters as cell types (F). Distribution of individual CRISPR/Cas9 perturbations across single cell transcriptomes. Note the offset position of Foxf^CRISPR^ transcriptomes (purple).

**Figure 5-S2.**
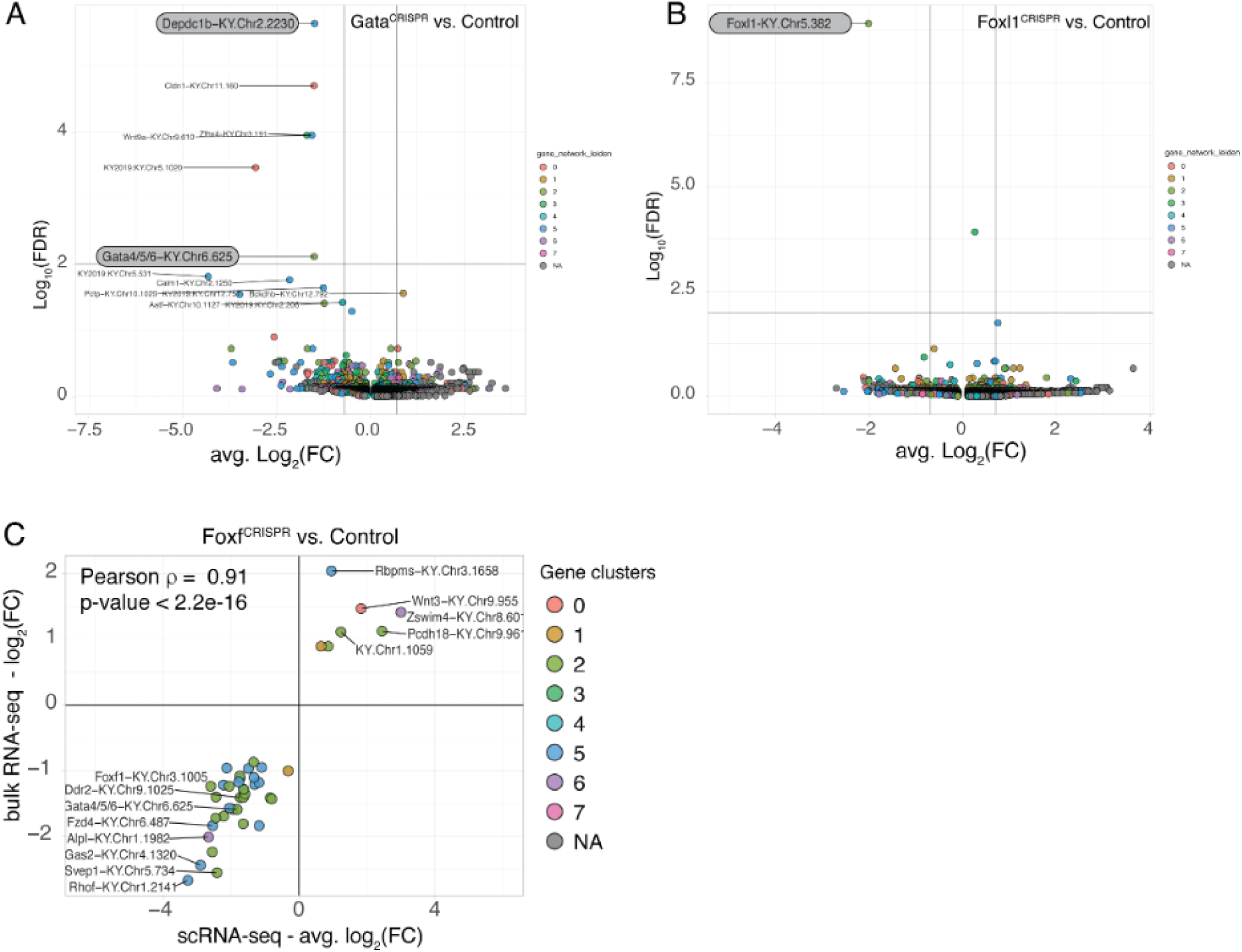
Results of the CRISPR mutagenesis targeting the Gata4/5/6, Foxl1, and Foxf loci followed by scRNA-seq of the cardiogenic and cardiopharyngeal lineages. **A.** Volcano plot of log2 fold gene expression change in Gata4/5/6 vs Control conditions with colors of genes corresponding to their Leiden cluster identity. **B**. Volcano plot of log2 fold gene expression change in Foxl1 vs Control conditions. For both A and B the Leiden clusters shown correspond to the Leiden clusters shown in Figure 2I. **C.** Log2 fold-change for top DE genes in Foxf^CRISPR^ vs controls comparing the scRNA-seq from this paper to the bulk RNA-seq of FACS-sorted cells from Racioppi et al 2019, with significant correlation values indicated.

**Figure 6-S1.**
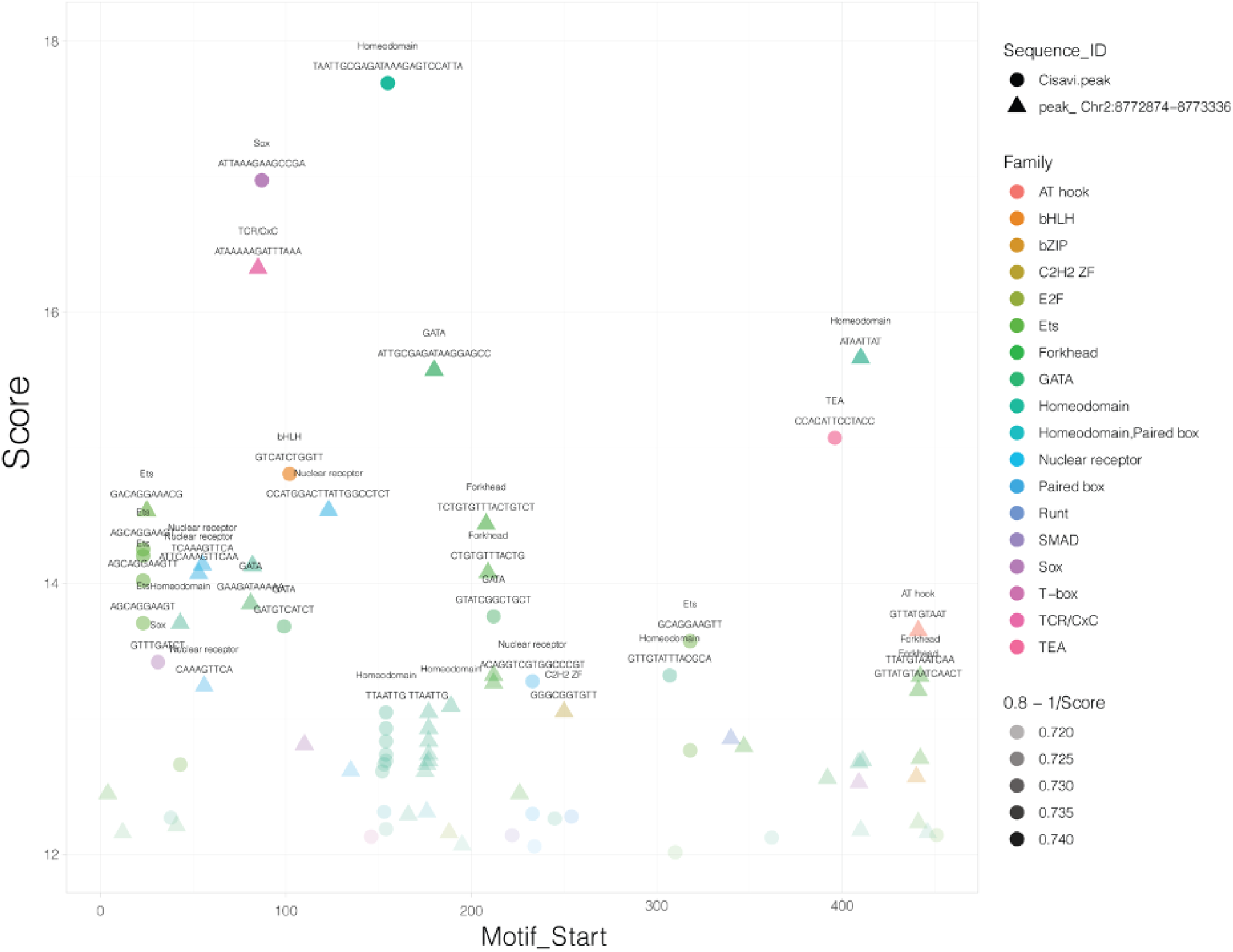
Putative Transcription Factor Binding Motifs in the *Depdc1b* enhancer conserved between *Ciona robusta* and *Ciona savignyi*. The sequences used correspond to the alignment showed in Figure 4, and the x axis correspond to the position on each sequence. The y axis shows the motif scores obtained from Cis-BP, using motifs inferred by homology for Ciona homologs of transcription factors indicated as color codes. Transparency is also inverse correlated with motif score. The shapes correspond to the species, Ciona savignyi (dots) or Ciona robusta (triangles).

## Notes

### Competing Interest Statement

The authors have declared no competing interest.

## References

1. Aviv, R., Teichmann, S.A., Lander, E.S., Ido, A., Christophe, B., Ewan, B., Bernd, B., Campbell, P., Piero, C., Menna, C., et al. (2017). The Human Cell Atlas. eLife; Cambridge 6. 10.7554/eLife.27041.

2. Cao, J., Spielmann, M., Qiu, X., Huang, X., Ibrahim, D.M., Hill, A.J., Zhang, F., Mundlos, S., Christiansen, L., Steemers, F.J., et al. (2019). The single-cell transcriptional landscape of mammalian organogenesis. Nature 566, 496–502.

3. Levy, S., Elek, A., Grau-Bové, X., Menéndez-Bravo, S., Iglesias, M., Tanay, A., Mass, T., and Sebé-Pedrós, A. (2021). A stony coral cell atlas illuminates the molecular and cellular basis of coral symbiosis, calcification, and immunity. Cell 184, 2973–2987.e18.

4. Moris, N., Pina, C., and Arias, A.M. (2016). Transition states and cell fate decisions in epigenetic landscapes. Nat. Rev. Genet. 17, 693–703.

5. Levine, M., and Tjian, R. (2003). Transcription regulation and animal diversity. Nature 424, 147–151.

6. Levine, M., and Davidson, E.H. (2005). Gene regulatory networks for development. Proc. Natl. Acad. Sci. U. S. A. 102, 4936–4942.

7. Harashima, H., Dissmeyer, N., and Schnittger, A. (2013). Cell cycle control across the eukaryotic kingdom. Trends Cell Biol. 23, 345–356.

8. Pagano, M. (2013). Cell Cycle Control (Springer Berlin Heidelberg).

9. Nurse, P. (2000). A long twentieth century of the cell cycle and beyond. Cell 100, 71–78.

10. Nair, G., Walton, T., Murray, J.I., and Raj, A. (2013). Gene transcription is coordinated with, but not dependent on, cell divisions during C. elegans embryonic fate specification. Development 140, 3385–3394.

11. Kukreja, K., Patel, N., Megason, S.G., and Klein, A.M. (2023). Global decoupling of cell differentiation from cell division in early embryo development. bioRxiv. 10.1101/2023.07.29.551123.

12. Hao, Y., Stuart, T., Kowalski, M.H., Choudhary, S., Hoffman, P., Hartman, A., Srivastava, A., Molla, G., Madad, S., Fernandez-Granda, C., et al. (2024). Dictionary learning for integrative, multimodal and scalable single-cell analysis. Nat. Biotechnol. 42, 293–304.

13. Wolf, F.A., Angerer, P., and Theis, F.J. (2018). SCANPY: large-scale single-cell gene expression data analysis. Genome Biol. 19, 15.

14. Pauklin, S., and Vallier, L. (2013). The cell-cycle state of stem cells determines cell fate propensity. Cell 155, 135–147.

15. Yiangou, L., Grandy, R.A., Osnato, A., Ortmann, D., Sinha, S., and Vallier, L. (2019). Cell cycle regulators control mesoderm specification in human pluripotent stem cells. J. Biol. Chem. 294, 17903–17914.

16. Soufi, A., and Dalton, S. (2016). Cycling through developmental decisions: how cell cycle dynamics control pluripotency, differentiation and reprogramming. Development 143, 4301–4311.

17. Dalton, S. (2015). Linking the Cell Cycle to Cell Fate Decisions. Trends Cell Biol. 25, 592–600.

18. Otsuki, L., and Brand, A.H. (2018). Cell cycle heterogeneity directs the timing of neural stem cell activation from quiescence. Science 360, 99–102.

19. El-Danaf, R.N., Rajesh, R., and Desplan, C. (2023). Temporal regulation of neural diversity in Drosophila and vertebrates. Semin. Cell Dev. Biol. 142, 13–22.

20. Doe, C.Q. (2017). Temporal Patterning in the Drosophila CNS. Annu. Rev. Cell Dev. Biol. 33, 219–240.

21. Edgar, B.A., Lehman, D.A., and O’Farrell, P.H. (1994). Transcriptional regulation of string (cdc25): a link between developmental programming and the cell cycle. Development 120, 3131–3143.

22. Edgar, B.A., and O’Farrell, P.H. (1989). Genetic control of cell division patterns in the Drosophila embryo. Cell 57, 177–187.

23. Di Talia, S., and Wieschaus, E.F. (2012). Short-term integration of Cdc25 dynamics controls mitotic entry during Drosophila gastrulation. Dev. Cell 22, 763–774.

24. Momen-Roknabadi, A., Di Talia, S., and Wieschaus, E. (2016). Transcriptional Timers Regulating Mitosis in Early Drosophila Embryos. Cell Rep. 16, 2793–2801.

25. Ogura, Y., and Sasakura, Y. (2016). Developmental Control of Cell-Cycle Compensation Provides a Switch for Patterned Mitosis at the Onset of Chordate Neurulation. Dev. Cell 37, 148–161.

26. Dumollard, R., Hebras, C., Besnardeau, L., and McDougall, A. (2013). Beta-catenin patterns the cell cycle during maternal-to-zygotic transition in urochordate embryos. Dev. Biol. 384, 331–342.

27. Kobayashi, K., Tokuoka, M., Sato, H., Ariyoshi, M., Kawahara, S., Fujiwara, S., Kishimoto, T., and Satou, Y. (2022). Regulators specifying cell fate activate cell cycle regulator genes to determine cell numbers in ascidian larval tissues. Development 149. 10.1242/dev.201218.

28. Guner-Ataman, B., González-Rosa, J.M., Shah, H.N., Butty, V.L., Jeffrey, S., Abrial, M., Boyer, L.A., Burns, C.G., and Burns, C.E. (2018). Failed Progenitor Specification Underlies the Cardiopharyngeal Phenotypes in a Zebrafish Model of 22q11.2 Deletion Syndrome. Cell Rep. 24, 1342–1354.e5.

29. Cirino, A., Aurigemma, I., Franzese, M., Lania, G., Righelli, D., Ferrentino, R., Illingworth, E., Angelini, C., and Baldini, A. (2020). Chromatin and Transcriptional Response to Loss of TBX1 in Early Differentiation of Mouse Cells. Front Cell Dev Biol 8, 571501.

30. Swedlund, B., and Lescroart, F. (2020). Cardiopharyngeal Progenitor Specification: Multiple Roads to the Heart and Head Muscles. Cold Spring Harb. Perspect. Biol. 12. 10.1101/cshperspect.a036731.

31. Rana, M.S., Théveniau-Ruissy, M., De Bono, C., Mesbah, K., Francou, A., Rammah, M., Domínguez, J.N., Roux, M., Laforest, B., Anderson, R.H., et al. (2014). Tbx1 coordinates addition of posterior second heart field progenitor cells to the arterial and venous poles of the heart. Circ. Res. 115, 790–799.

32. Nomaru, H., Liu, Y., De Bono, C., Righelli, D., Cirino, A., Wang, W., Song, H., Racedo, S.E., Dantas, A.G., Zhang, L., et al. (2021). Single cell multi-omic analysis identifies a Tbx1-dependent multilineage primed population in murine cardiopharyngeal mesoderm. Nat. Commun. 12, 6645.

33. Adachi, N., Bilio, M., Baldini, A., and Kelly, R.G. (2020). Cardiopharyngeal mesoderm origins of musculoskeletal and connective tissues in the mammalian pharynx. Development 147. 10.1242/dev.185256.

34. Kaplan, N., Razy-Krajka, F., and Christiaen, L. (2015). Regulation and evolution of cardiopharyngeal cell identity and behavior: insights from simple chordates. Curr. Opin. Genet. Dev. 32, 119–128.

35. Wang, W., Niu, X., Stuart, T., Jullian, E., Mauck, W.M., Kelly, R.G., Satija, R., and Christiaen, L. (2019). A single-cell transcriptional roadmap for cardiopharyngeal fate diversification. Preprint, 10.1038/s41556-019-0336-z 10.1038/s41556-019-0336-z.

36. Wang, W., Razy-Krajka, F., Siu, E., Ketcham, A., and Christiaen, L. (2013). NK4 antagonizes Tbx1/10 to promote cardiac versus pharyngeal muscle fate in the ascidian second heart field. PLoS Biol. 11, e1001725.

37. Razy-Krajka, F., Gravez, B., Kaplan, N., Racioppi, C., Wang, W., and Christiaen, L. (2018). An FGF-driven feed-forward circuit patterns the cardiopharyngeal mesoderm in space and time. Elife 7. 10.7554/eLife.29656.

38. Tolkin, T., and Christiaen, L. (2016). Rewiring of an ancestral Tbx1/10-Ebf-Mrf network for pharyngeal muscle specification in distinct embryonic lineages. Development 143, 3852–3862.

39. Song, M., Yuan, X., Racioppi, C., Leslie, M., Stutt, N., Aleksandrova, A., Christiaen, L., Wilson, M.D., and Scott, I.C. (2022). GATA4/5/6 family transcription factors are conserved determinants of cardiac versus pharyngeal mesoderm fate. Sci Adv 8, eabg0834.

40. Gline, S., Kaplan, N., Bernadskaya, Y., Abdu, Y., and Christiaen, L. (2015). Surrounding tissues canalize motile cardiopharyngeal progenitors towards collective polarity and directed migration. Preprint, 10.1242/dev.115444 10.1242/dev.115444.

41. Beh, J., Shi, W., Levine, M., Davidson, B., and Christiaen, L. (2007). FoxF is essential for FGF-induced migration of heart progenitor cells in the ascidian Ciona intestinalis. Development 134, 3297–3305.

42. Donzelli, M., and Draetta, G.F. (2003). Regulating mammalian checkpoints through Cdc25 inactivation. EMBO Rep. 4, 671–677.

43. Hotta, K., Mitsuhara, K., Takahashi, H., Inaba, K., Oka, K., Gojobori, T., and Ikeo, K. (2007). A web-based interactive developmental table for the ascidian Ciona intestinalis, including 3D real-image embryo reconstructions: I. From fertilized egg to hatching larva. Dev. Dyn. 236, 1790–1805.

44. Schönenberger, F., Deutzmann, A., Ferrando-May, E., and Merhof, D. (2015). Discrimination of cell cycle phases in PCNA-immunolabeled cells. BMC Bioinformatics 16, 180.

45. Tjärnberg, A., Mahmood, O., Jackson, C.A., Saldi, G.-A., Cho, K., Christiaen, L.A., and Bonneau, R.A. Optimal tuning of weighted kNN- and diffusion-based methods for denoising single cell genomics data. Preprint, 10.1101/2020.02.28.970202 10.1101/2020.02.28.970202.

46. Christiaen, L., Davidson, B., Kawashima, T., Powell, W., Nolla, H., Vranizan, K., and Levine, M. (2008). The transcription/migration interface in heart precursors of Ciona intestinalis. Science 320, 1349–1352.

47. Ohta, N., and Christiaen, L. (2021). Cellular remodeling and JAK inhibition promote zygotic gene expression in the*Ciona*germline. bioRxiv. 10.1101/2021.07.12.452040.

48. Cao, C., Lemaire, L.A., Wang, W., Yoon, P.H., Choi, Y.A., Parsons, L.R., Matese, J.C., Wang, W., Levine, M., and Chen, K. (2019). Comprehensive single-cell transcriptome lineages of a proto-vertebrate. Nature 571, 349–354.

49. Stolfi, A., Gandhi, S., Salek, F., and Christiaen, L. (2014). Tissue-specific genome editing in Ciona embryos by CRISPR/Cas9. Development 141, 4115–4120.

50. Gandhi, S., Haeussler, M., Razy-Krajka, F., Christiaen, L., and Stolfi, A. (2017). Evaluation and rational design of guide RNAs for efficient CRISPR/Cas9-mediated mutagenesis in Ciona. Dev. Biol. 425, 8–20.

51. Rose, L.S., and Kemphues, K. (1998). The let-99 gene is required for proper spindle orientation during cleavage of the C. elegans embryo. Development 125, 1337–1346.

52. Tsou, M.-F.B., Hayashi, A., DeBella, L.R., McGrath, G., and Rose, L.S. (2002). LET-99 determines spindle position and is asymmetrically enriched in response to PAR polarity cues in C. elegans embryos. Development 129, 4469–4481.

53. Bernadskaya, Y.Y., Brahmbhatt, S., Gline, S.E., Wang, W., and Christiaen, L. (2019). Discoidin-domain receptor coordinates cell-matrix adhesion and collective polarity in migratory cardiopharyngeal progenitors. Preprint, 10.1038/s41467-018-07976-3 10.1038/s41467-018-07976-3.

54. Racioppi, C., Wiechecki, K.A., and Christiaen, L. (2019). Combinatorial chromatin dynamics foster accurate cardiopharyngeal fate choices. Elife 8. 10.7554/eLife.49921.

55. Woznica, A., Haeussler, M., Starobinska, E., Jemmett, J., Li, Y., Mount, D., and Davidson, B. (2012). Initial deployment of the cardiogenic gene regulatory network in the basal chordate, Ciona intestinalis. Dev. Biol. 368, 127–139.

56. Davidson, E.H. (2009). Network design principles from the sea urchin embryo. Curr. Opin. Genet. Dev. 19, 535–540.

57. Davidson, E.H. (2010). Emerging properties of animal gene regulatory networks. Nature 468, 911–920.

58. Lambert, S.A., Yang, A.W.H., Sasse, A., Cowley, G., Albu, M., Caddick, M.X., Morris, Q.D., Weirauch, M.T., and Hughes, T.R. (2019). Similarity regression predicts evolution of transcription factor sequence specificity. Nat. Genet. 51, 981–989.

59. Qiu, C., Cao, J., Martin, B.K., Li, T., Welsh, I.C., Srivatsan, S., Huang, X., Calderon, D., Noble, W.S., Disteche, C.M., et al. (2022). Systematic reconstruction of cellular trajectories across mouse embryogenesis. Nat. Genet. 54, 328–341.

60. Farrell, J.A., Wang, Y., Riesenfeld, S.J., Shekhar, K., Regev, A., and Schier, A.F. (2018). Single-cell reconstruction of developmental trajectories during zebrafish embryogenesis. Science 360. 10.1126/science.aar3131.

61. Briggs, J.A., Weinreb, C., Wagner, D.E., Megason, S., Peshkin, L., Kirschner, M.W., and Klein, A.M. (2018). The dynamics of gene expression in vertebrate embryogenesis at single-cell resolution. Science 360. 10.1126/science.aar5780.

62. Pauklin, S., Madrigal, P., Bertero, A., and Vallier, L. (2016). Initiation of stem cell differentiation involves cell cycle-dependent regulation of developmental genes by Cyclin D. Genes Dev. 30, 421–433.

63. Dalton, S. (2013). G1 compartmentalization and cell fate coordination. Cell 155, 13–14.

64. Gonzales, K.A.U., Liang, H., Lim, Y.-S., Chan, Y.-S., Yeo, J.-C., Tan, C.-P., Gao, B., Le, B., Tan, Z.-Y., Low, K.-Y., et al. (2015). Deterministic Restriction on Pluripotent State Dissolution by Cell-Cycle Pathways. Cell 162, 564–579.

65. Guignard, L., Fiúza, U.-M., Leggio, B., Laussu, J., Faure, E., Michelin, G., Biasuz, K., Hufnagel, L., Malandain, G., Godin, C., et al. (2020). Contact area-dependent cell communication and the morphological invariance of ascidian embryogenesis. Science 369. 10.1126/science.aar5663.

66. Lemaire, P. (2009). Unfolding a chordate developmental program, one cell at a time: invariant cell lineages, short-range inductions and evolutionary plasticity in ascidians. Dev. Biol. 332, 48–60.

67. Sun, J., Durmaz, A.D., Babu, A., Macabenta, F., and Stathopoulos, A. (2024). Two sequential gene expression programs bridged by cell division support long-distance collective cell migration. Development 151. 10.1242/dev.202262.

68. Chen, J., Cheung, F., Shi, R., Zhou, H., Lu, W., and CHI Consortium (2018). PBMC fixation and processing for Chromium single-cell RNA sequencing. J. Transl. Med. 16, 198.

69. Dardaillon, J., Dauga, D., Simion, P., Faure, E., Onuma, T.A., DeBiasse, M.B., Louis, A., Nitta, K.R., Naville, M., Besnardeau, L., et al. (2020). ANISEED 2019: 4D exploration of genetic data for an extended range of tunicates. Nucleic Acids Res. 48, D668–D675.

70. Heinz, S., Benner, C., Spann, N., Bertolino, E., Lin, Y.C., Laslo, P., Cheng, J.X., Murre, C., Singh, H., and Glass, C.K. (2010). Simple combinations of lineage-determining transcription factors prime cis-regulatory elements required for macrophage and B cell identities. Mol. Cell 38, 576–589.

